# Shaped by meaning, weighted by reliability: New insights into multisensory integration

**DOI:** 10.64898/2026.01.05.697596

**Authors:** Elizaveta Sycheva, Léa St-Gelais, Karim Jerbi, Shahab Bakhtiari, Franco Lepore, Vanessa Hadid

## Abstract

In everyday perception, sensory information is rarely interpreted in isolation. Instead, observers generate expectations about what is likely to occur next, using contextual information to guide decisions, particularly when sensory input is weak, ambiguous, or uncertain. These expectations can facilitate behaviour when they align with incoming evidence, but impose costs when they conflict, shaping how information from different senses contributes to behaviour. However, it remains unclear what kind of multisensory process is engaged when meaning guides decisions, and whether multisensory facilitation and non-independence reflect a single integrative mechanism or distinct routes shaped by context. Here, we examined how semantic context, visual clarity, and the relationship between auditory and visual signals jointly shape behaviour in a semantic correspondence task using dynamic, naturalistic events. Participants judged whether a target matched a preceding concept prime, establishing an expectation at the level of meaning. Targets were auditory-only, visual-only (clear or blurry), or audiovisual (AV), with AV components conveying either the same or different meanings across modalities. AV facilitation was not a general consequence of stimulus redundancy. When the prime accurately predicted the target and auditory and visual signals conveyed the same meaning, degrading visual input increased the contribution of auditory information, leading to faster responses and violations of the race-model inequality. By contrast, when the prime indicated the target but auditory and visual signals conveyed competing meanings, AV stimulation produced robust behavioural costs relative to vision. When the prime did not match the target, congruent and incongruent audiovisual events produced comparable effects, and race-model violations were observed even when auditory and visual signals encoded different concepts, indicating convergence at the level of the response rather than shared perceptual content. Finally, exploratory clustering revealed two participant subgroups with comparable overall response speed but systematically different multisensory profiles when semantic expectations did not allow reliable anticipation of the upcoming event. Together, these findings show that multisensory effects in semantic decisions are shaped by meaning and arise through two functionally distinct routes: one in which reduced sensory reliability promotes joint use of congruent audiovisual information, and another in which perceptually different signals converge on the same response when semantic expectations are not met.

**Highlights:**

1. Semantic expectations shape how multisensory information is used to guide responses.
2. Reduced sensory reliability enables multisensory facilitation when signals convey the same meaning.
3. Multisensory effects arise when sensory signals support the same response.
4. Individuals adopt distinct strategies for resolving uncertainty in multisensory decisions.

## Main

Everyday perception unfolds in a predictive context. Rather than interpreting sensory information in isolation, observers continuously generate expectations about what is likely to occur next, using contextual information to guide decisions, particularly when sensory input is weak, ambiguous, or uncertain (Friston, 2005, 2010; Bar, 2007; Clark, 2013). These expectations can facilitate behaviour when they align with incoming evidence, but impose measurable costs when they are violated. Crucially, predictions also shape how information from different senses is evaluated and combined during decision-making (de Lange et al., 2018).

Multisensory input plays a central role in this process. By combining information across modalities, observers can often respond more robustly and efficiently, improving detection, discrimination, and recognition (Ernst & Bülthoff, 2004; Stein & Stanford, 2007; Chandrasekaran, 2017; Otto et al., 2013). AV signals are especially relevant in natural environments, where sounds and visual events frequently co-occur and can jointly support behaviour (Gao et al., 2023). However, multisensory benefits are not uniform. Depending on sensory reliability and contextual factors, combining information can facilitate decisions, add little beyond a single modality, or even incur behavioural costs. Understanding when and why multisensory information benefits decision-making in naturalistic contexts remains a central challenge.

Previous work has shown that when one sensory modality provides weak or degraded evidence, information from another modality can exert a stronger influence on behaviour, often described in terms of reliability-based multisensory facilitation or inverse effectiveness (Ernst & Bülthoff, 2004; Stein & Meredith, 1993; Holmes, 2007). However, these frameworks are typically grounded in perceptual decision tasks with well-defined stimulus–response mappings and do not directly address how sensory information is recruited when decisions depend on semantic interpretation and explicit correspondence judgments. When decisions depend on meaning rather than detection or discrimination, the contribution of each sensory modality depends critically on how efficiently it supports the required judgement (Otto et al., 2013; Innes & Otto, 2019). Accordingly, whether multisensory input provides a behavioural advantage depends on how effectively each modality supports access to the relevant meaning. When one modality already supports efficient decisions, adding information from another sense may yield little benefit relative to that modality, even if it improves performance compared to a less efficient channel.

Beyond sensory reliability, top-down contextual predictions influence how incoming evidence is interpreted and prioritised. In semantic priming paradigms, such expectations are understood to operate at the level of conceptual representations, increasing the accessibility of task-relevant meanings and biasing how subsequent sensory evidence is interpreted and evaluated (Bar, 2007; de Lange et al., 2018). Accordingly, the prime does not specify the perceptual form of the upcoming event, but establishes a semantic hypothesis that must be evaluated against incoming sensory information. Prior knowledge, learned associations, and semantic context can further bias these expectations, facilitating decisions when predictions are confirmed and imposing costs when they are violated (Doehrmann & Naumer, 2008; Talsma et al., 2010; Gau & Noppeney, 2016; de Lange et al., 2018). In naturalistic settings, such predictions often interact with the correspondence between concurrently available sensory signals. When auditory and visual information support the same interpretation, they can reinforce one another; when they convey competing information, they may introduce conflict during response generation. Although both contextual expectations and cross-modal correspondence have been shown to influence multisensory behaviour (Chen & Spence, 2010, 2011; Marini & Maravita, 2017), they are typically examined in isolation. Their joint contributions to multisensory facilitation and cost, particularly in dynamic and semantically meaningful contexts, remain poorly understood. Under these conditions, it remains unclear whether similar multisensory behavioural outcomes reflect a common underlying process, or instead arise from distinct strategies shaped by different interactions between sensory reliability, contextual predictions, and cross-modal correspondence.

Interpreting multisensory facilitation also requires careful consideration of behavioural benchmarks. Faster responses to AV stimuli are often described as a redundant signals effect, namely improved response speed when two signals are present compared with either alone (Miller, 1982; Otto & Mamassian, 2012; Innes & Otto, 2019). However, such speed-ups do not necessarily reflect multisensory interaction. Critically, it remains unclear when AV responses exceed what probability summation alone can explain, and whether such departures from independence arise systematically from interactions between sensory reliability and contextual predictions.

In the race model (Miller, 1982), auditory and visual signals are processed independently and in parallel, with the faster channel triggering the response on each trial. Under this assumption, AV response times are constrained by an upper limit known as the Miller bound, which reflects the maximum facilitation expected from probability summation alone. When observed AV responses exceed this bound, the race-model inequality is violated, indicating that auditory and visual information cannot be fully independent and must jointly contribute before the response is generated (Chua et al., 2022). Such race-model violations therefore provide evidence for decision-level multisensory contributions, without implying perceptual fusion or a specific neural integration mechanism (Diederich & Colonius, 2004; Colonius & Diederich, 2017).

Finally, it remains unclear whether observers adapt to sensory and contextual constraints in uniform ways. Substantial individual variability in multisensory behaviour has been documented, yet it is often treated as noise rather than as a meaningful feature of perceptual decision-making (Mahoney et al., 2011; Magnotti & Beauchamp, 2018). Challenging contexts, such as reduced sensory reliability or violated expectations, may reveal divergent strategies for exploiting available information. Whether such variability reflects arbitrary differences or systematic adaptations to decision uncertainty remains an open question.

The present study addresses these issues by examining when AV information benefits behavioural decisions, and for whom these benefits emerge, using dynamic and semantically meaningful AV events. We manipulated bottom-up sensory reliability by varying visual clarity, top-down contextual predictions by manipulating prime–target coherence, and cross-modal correspondence by varying AV congruency. Using accuracy, reaction times, response-time distributions, race-model analyses, and descriptive weighting measures, we quantified multisensory benefits and costs across contexts. We further assessed whether participants exhibited distinct behavioural profiles in how they adapt to these constraints, providing insight into variability in multisensory decision strategies under uncertainty.

## Results

Participants (*n* = 42) completed a semantic correspondence task in which spoken primes were followed by naturalistic AV stimuli drawn from five semantic categories. Across four blocks, 117 stimuli varied systematically in modality, visual clarity, semantic coherence, and repetition, yielding seven conditions spanning unimodal and AV trials. Participants indicated whether each stimulus matched the preceding prime, allowing us to characterize how these factors shape multisensory semantic decisions (Fig. 1).

**Figure 1.**
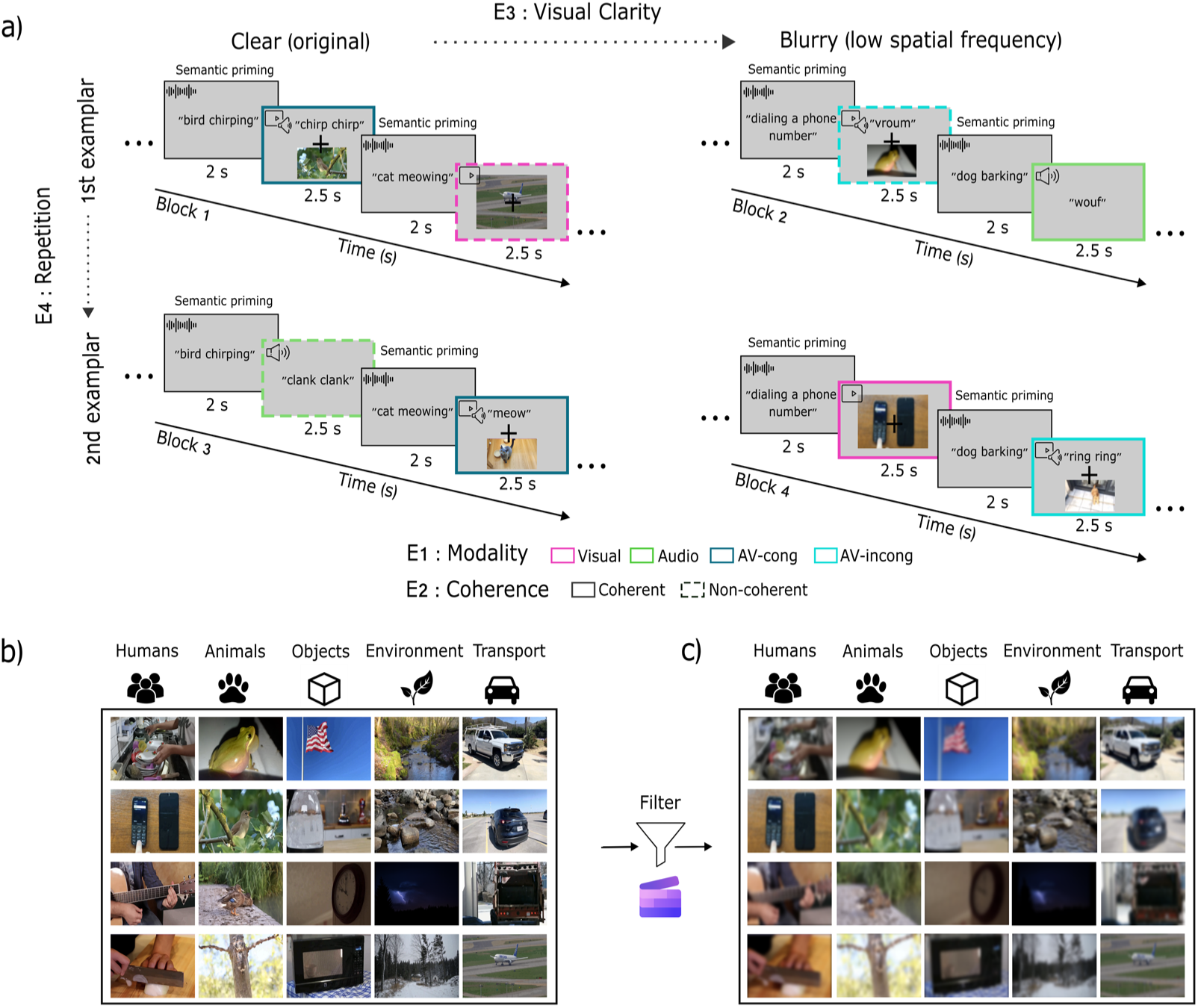
Task design and experimental conditions. (a) Trial structure. Each trial began with a 2.5-s spoken auditory prime naming a semantic concept, followed by a 2-s target stimulus presented in one of seven conditions: auditory-only, visual-only (clear), visual-only (blurry), AV congruent (clear), AV congruent (blurry), AV incongruent (clear), or AV incongruent (blurry). Participants judged whether the target matched the preceding prime (“1” = match, “2” = mismatch). (b) Example clear visual stimuli drawn from the five semantic categories (humans, animals, objects, environment, transport). (c) Blurred versions of the same example stimuli shown in (b), illustrating the manipulation of visual clarity. Blurring selectively reduced high spatial-frequency information while preserving global scene structure; auditory signals were unchanged.

**Figure 2.**
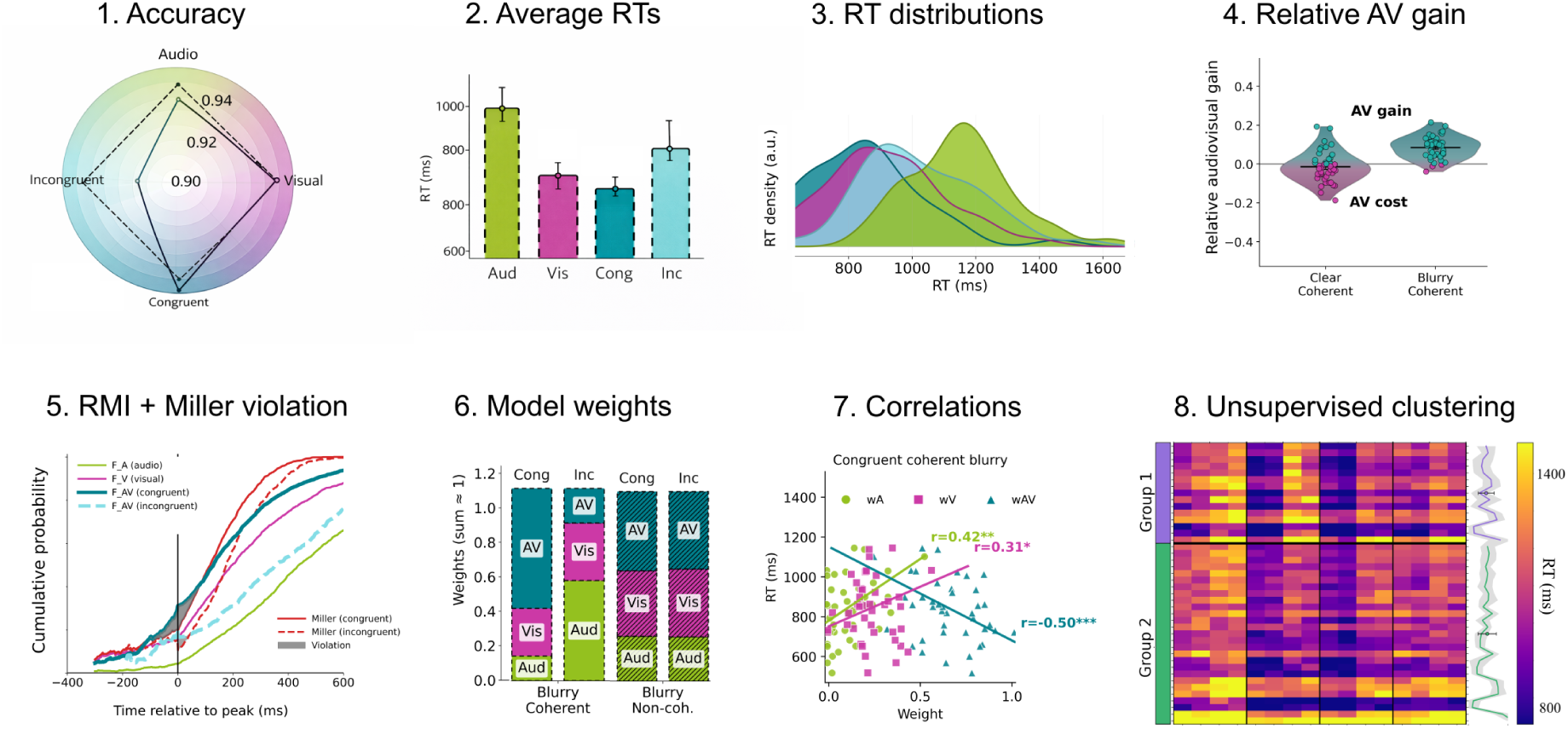
Overview of behavioural analyses. Schematic overview of the analytic pipeline. Behavioural analyses included (1) accuracy and (2) RTs, (3) full RT distributions, (4) AV gains and costs relative to unimodal baselines, (5) race-model inequality (Miller bound) metrics, (6) descriptive auditory, visual, and AV weighting indices, and their (7) associations with RTs. (8) RTs were further entered into an exploratory, data-driven clustering analysis to characterize variability in multisensory response profiles across participants.

### Behavioural accuracy: dominant effects of modality and prime–target coherence Overall performance and main effects

Participants performed the task with high accuracy overall (mean = 0.93), well above chance. Accuracy varied significantly with modality, semantic coherence, and repetition (all *p* < 0.01), with modality producing the strongest effect (Fig. 3a–c). Accuracy was highest for AV-congruent trials (0.97) and lowest for AV-incongruent trials (0.85), with visual-only (0.96) and auditory-only (0.93) performance intermediate (*p* < 0.001; Fig. 3a). Collapsing across modality and clarity, semantic coherence improved accuracy overall (prime–target match: 0.95 vs. mismatch: 0.91; *p* < 0.01; Fig. 3a). Repetition increased accuracy from first to second exemplar (0.92 to 0.95; *p* < 0.01; Fig. 3c), whereas visual clarity did not yield a significant main effect when averaged across modality and semantic context (clear: 0.96 vs. blurry: 0.94; *p* = 0.78; Fig. 3b).

**Figure 3.**
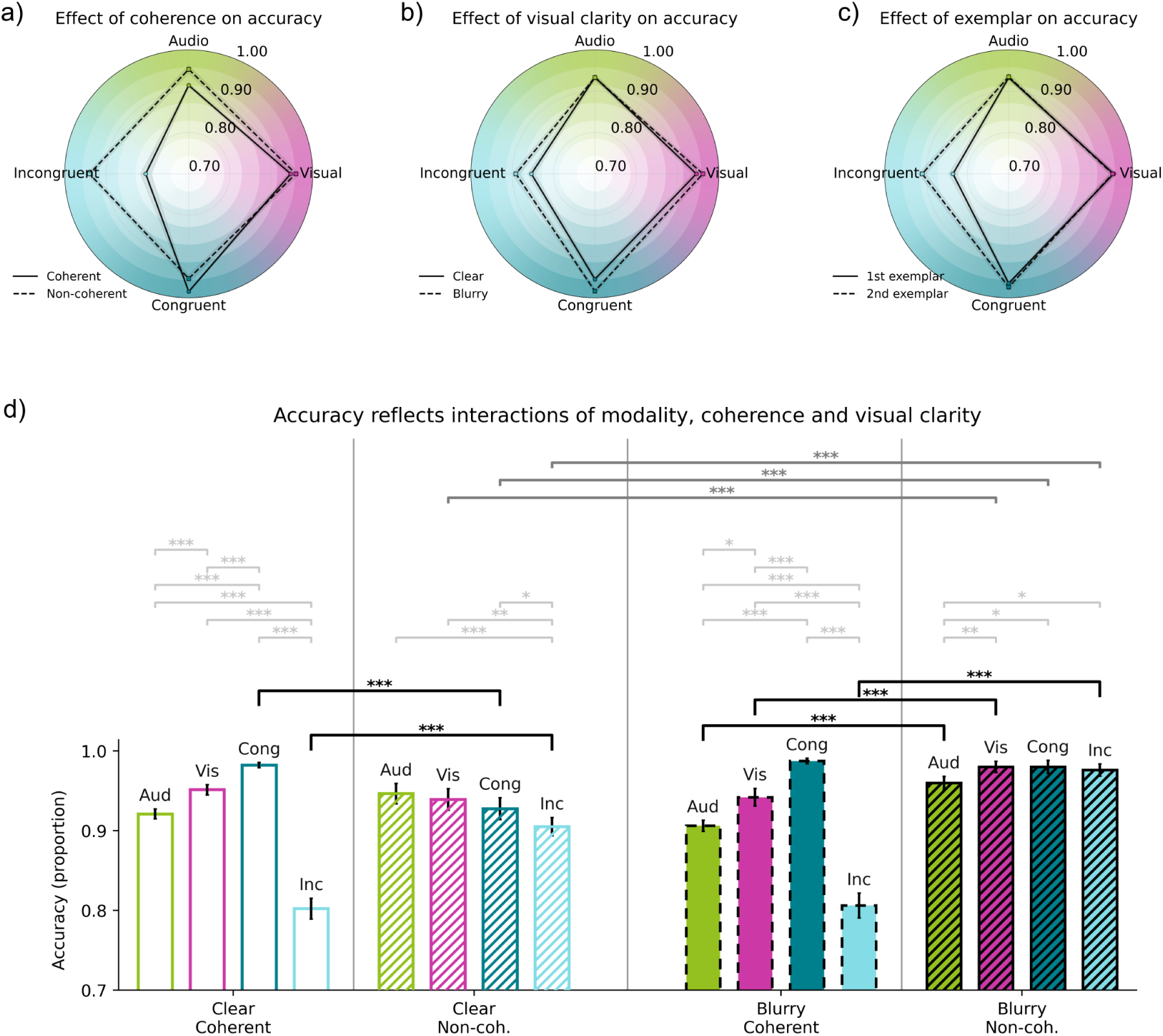
Accuracy varies with modality, coherence, and clarity. (a–c) Radar plots show mean accuracy (proportion correct) across response modalities (auditory, visual, AV-congruent, AV-incongruent), highlighting the effect of a single factor while collapsing across the others. (a) Effect of semantic coherence (coherent vs. non-coherent). (b) Effect of visual clarity (clear vs. blurry). (c) Effect of exemplar repetition (first vs. second exemplar). Solid and dashed lines denote contrasted levels within each panel. (d) Bar plots show mean ± SEM accuracy for the full interaction between modality, semantic coherence, and visual clarity, separated by coherence (coherent vs. non-coherent) and clarity (clear vs. blurry). Hatched bars indicate non-coherent conditions. Horizontal brackets denote within-subject contrasts surviving BH–FDR correction (*p* < 0.05, \**p* < 0.01, \*\**p* < 0.001). Across panels, AV-congruent trials yielded the highest accuracy, whereas AV-incongruent trials showed pronounced accuracy costs under semantic coherence, with reduced incongruence penalties under prime–target mismatch and blurry visual input.

### Interactions among modality, coherence, and clarity

A robust modality × semantic coherence interaction (*p* < 0.001) indicated that the behavioural cost of AV incongruence depended on semantic context (Supplementary Table 1). Under prime–target match, accuracy for AV-incongruent trials dropped markedly (0.80) relative to AV-congruent (0.98) and visual-only trials (0.94), whereas under prime–target mismatch AV-incongruent accuracy was substantially higher (0.94), indicating a reduced incongruence penalty when semantic expectations were not met (Fig. 3d).

A modality × visual clarity interaction (*p* < 0.05) further showed that clarity effects differed across modalities. In visual-only trials, accuracy was higher for clear than blurred stimuli under prime–target match (0.951 ± 0.006 vs. 0.942 ± 0.011), whereas the opposite pattern was observed under mismatch, with higher accuracy for blurred than clear stimuli (0.980 ± 0.007 vs. 0.939 ± 0.013; Fig. 3d). By contrast, accuracy for AV-congruent trials remained near ceiling across clarity levels and did not reliably depend on visual clarity (Fig. 3a,b). Given the overall ceiling-level accuracy, this crossover pattern likely reflects context-dependent decision dynamics rather than a general advantage of visual degradation. Repetition effects were modest and did not qualitatively alter the observed accuracy patterns across modality, semantic coherence, and visual clarity.

### Relative contribution of predictors

Variance decomposition confirmed that modality explained the largest share of unique variance in accuracy (partial *R²* = 0.039), followed by semantic coherence (0.018), repetition (0.010), the modality × coherence interaction (0.008), and visual clarity (0.005). Overall, modality and semantic coherence were the principal determinants of accuracy, with visual clarity and repetition exerting smaller, context-dependent influences (Supplementary Table 1).

### RTs: converging influences of modality, prime–target coherence, and clarity Overall RT patterns and main effects

Mean RTs were approximately 1005 ms and varied strongly with modality and semantic coherence, with additional context-dependent effects of clarity and repetition. Modality and coherence accounted for the largest differences.

Across modalities, auditory-only trials were slowest (1172 ms), followed by AV-incongruent (1081 ms), visual-only (1019 ms), and AV-congruent trials (980 ms). Auditory-only and AV-incongruent trials were significantly slower than visual-only trials (*p* < 1 × 10⁻³² and *p* < 1 × 10⁻¹⁶, respectively), whereas AV-congruent trials did not differ from visual-only trials (*p* = 0.82) (Fig. 4a).

**Figure 4.**
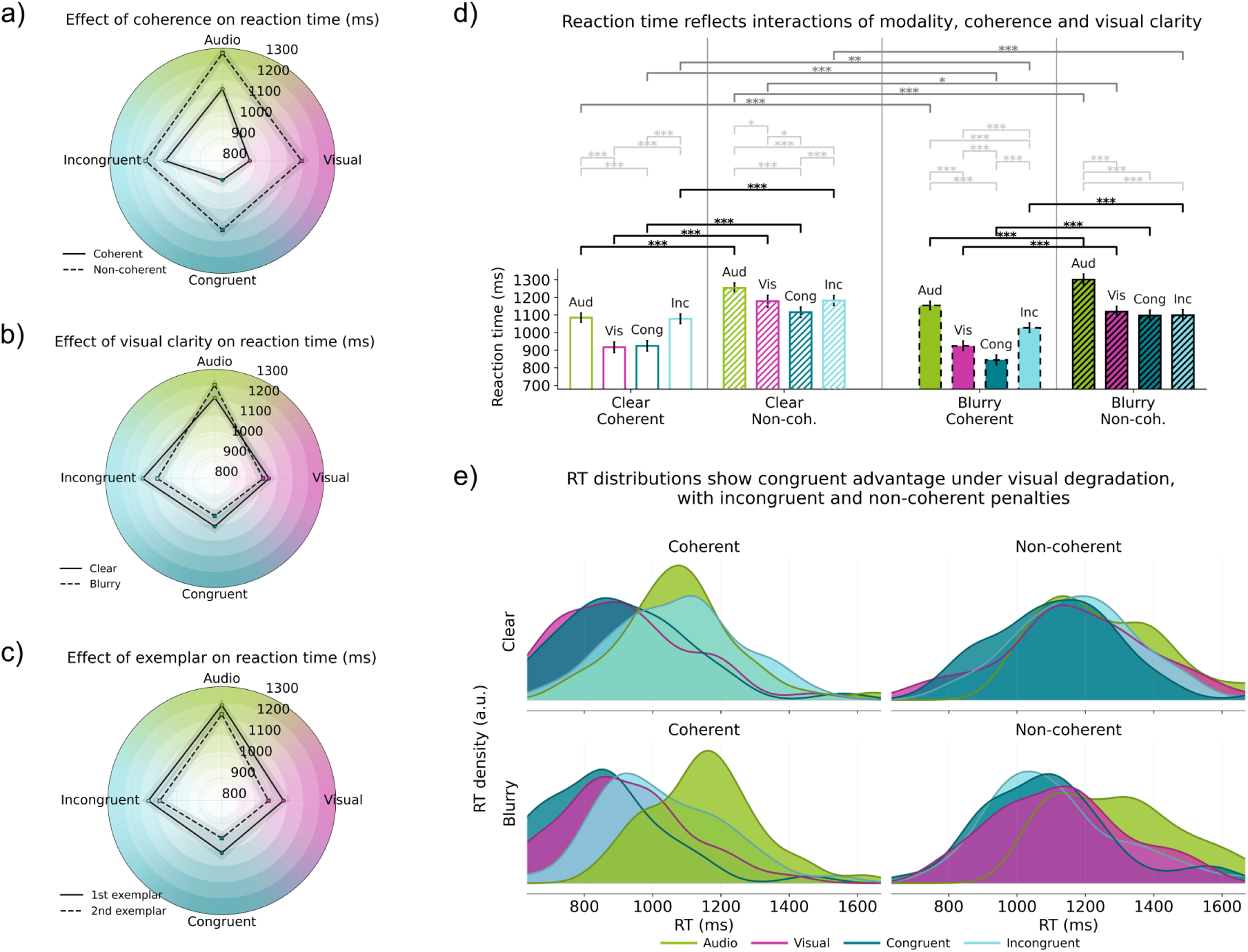
RTs vary with modality, coherence, and visual clarity. **(a)** Radar plots show mean ± SEM RTs across modalities (auditory, visual, AV-congruent, AV-incongruent) for each experimental contrast; solid and dashed lines denote contrasted levels (e.g., coherent vs. non-coherent, clear vs. blurry, first vs. second exemplar). **(b)** Condition-wise bar plots display mean ± SEM RTs for all modality × clarity × coherence combinations; hatched bars indicate non-coherent conditions. Horizontal brackets denote within-subject contrasts surviving BH–FDR correction (*p* < 0.05, *p* < 0.01, *p* < 0.001). **(c)** Ridgeline plots show full RT distributions by modality, clarity, and coherence, with earlier and narrower distributions for AV-congruent coherent conditions and broader, right-skewed distributions for auditory-only and non-coherent conditions.

Semantic non-coherence produced a similarly strong effect, with non-coherent trials approximately 175 ms slower than coherent trials (*p* < 1 × 10⁻²⁹). Visual clarity showed no reliable main effect (*p* = 0.58). Repetition produced modest speed-ups, with second exemplars faster than first exemplars across conditions (Fig. 4a).

### Interaction effects

A strong modality × coherence interaction (partial *R²* = 0.0087) indicated that coherence costs depended on modality (Supplementary Table 2). Non-coherent primes increased RTs across modalities, but coherence costs were significantly attenuated for auditory-only and AV-incongruent trials. Correspondingly, contrasts involving auditory × non-coherence (*p* = 5 × 10⁻⁶) and incongruent × non-coherence (*p* = 1 × 10⁻⁶) were reduced (Fig. 4b).

### Relative contribution of predictors

A modality × clarity interaction (partial *R²* = 0.0075) showed that auditory-only responses were slower under blurring, whereas visual-only and AV-congruent responses were slightly faster (Supplementary Table 2). A clarity × coherence interaction (*p* = 0.022) and a significant three-way interaction, most pronounced for AV-congruent trials (*p* = 5 × 10⁻⁴), further modulated RTs (Fig. 4b). Coherent, clear AV-congruent trials produced the fastest and most compact RT distributions, whereas non-coherent and auditory-only conditions yielded broader, right-skewed distributions extending beyond 1700–2000 ms. Repetition did not interact reliably with other factors (Fig. 4c).

Modality accounted for the largest unique share of RT variance (*R²* = 0.1017), followed by semantic coherence (0.0833), modality × coherence (0.0087), modality × clarity (0.0075), and clarity (0.0062). RTs, like accuracy, were therefore primarily driven by modality and semantic coherence at the group level, with visual clarity and related interactions modulating these effects in a context-dependent manner (Supplementary Table 2).

### Relative AV gain and cost relative to unimodal baselines AV performance relative to the auditory baseline

To characterize the magnitude and direction of multisensory facilitation, we quantified relative AV and costs with respect to unimodal auditory and visual baselines (Supplementary Table 3). Positive values indicate faster responses in the AV condition relative to the unimodal reference (AV gain), whereas negative values indicate slower AV responses (AV cost). This approach allows multisensory benefits and penalties to be interpreted separately with respect to each sensory stream (Fig. 5).

**Figure 5.**
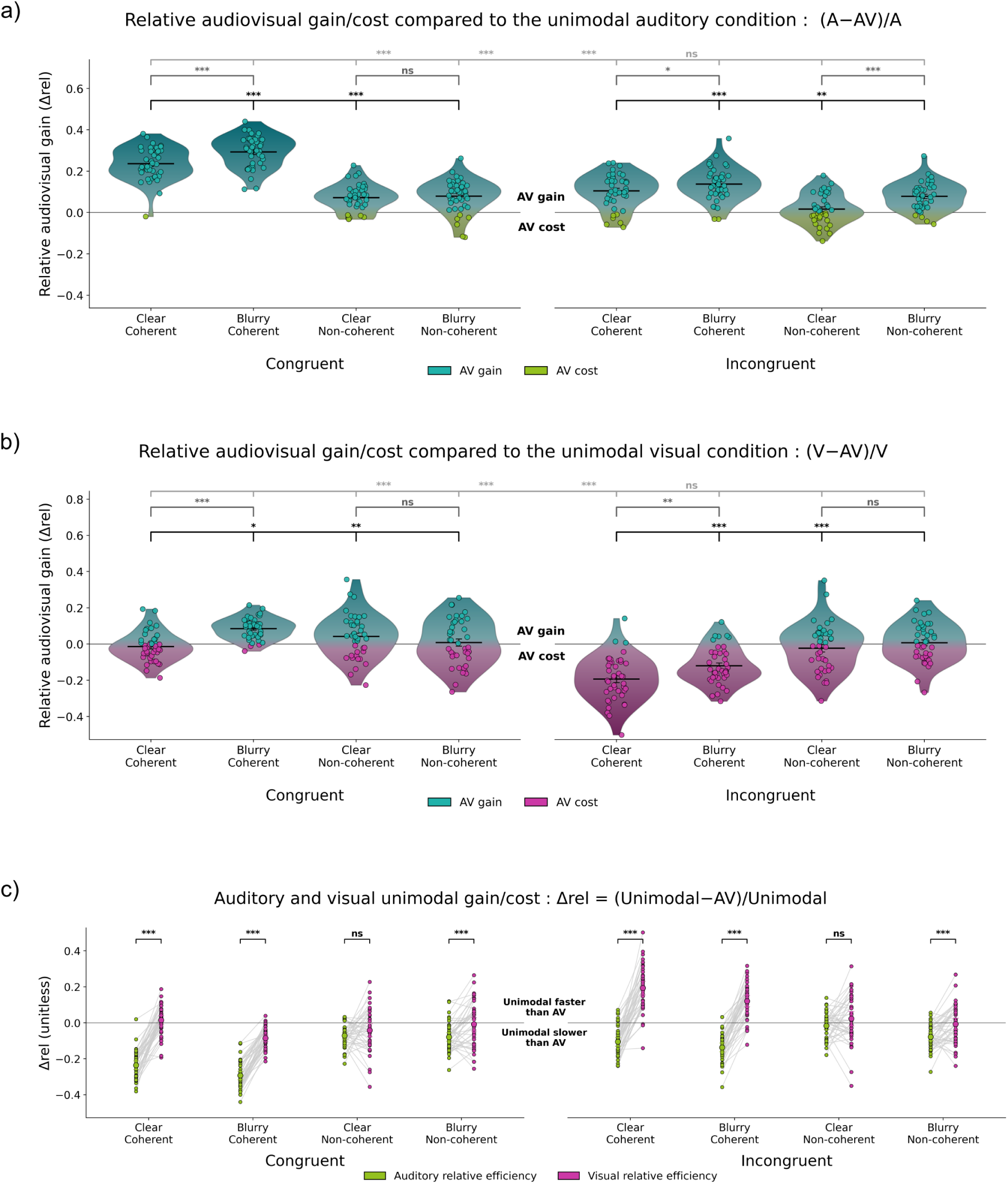
Relative AV gain and unimodal efficiency across clarity, coherence, and congruence. **(a)** Relative AV gain or cost computed using the auditory only condition as baseline. Positive values indicate faster AV responses relative to auditory only trials which correspond to AV gain and negative values indicate slower AV responses which correspond to AV cost. Violin plots show the distribution of relative differences Δrel across participants for each combination of visual clarity clear or blurry semantic coherence coherent or noncoherent with hatched fill and AV congruence congruent or incongruent. Individual participant values are shown with cyan dots indicating AV gains and green dots indicating AV costs and black horizontal bars indicate mean ± SEM. Horizontal brackets summarize paired comparisons between clear and blurry conditions congruent and incongruent AV conditions and coherent versus noncoherent contexts with significance levels corrected using BH FDR. **(b)** Same analysis as in (a) using the visual only condition as baseline. Positive values indicate faster AV responses relative to visual only trials and negative values indicate relative AV costs. Pink points denote AV costs relative to vision. **(c)** Relative unimodal efficiency indices for auditory and visual conditions across all clarity × coherence × congruence combinations. For each participant auditory indices shown in green and visual indices shown in magenta are connected by a line with larger circles indicating the mean ± 95% confidence interval. Negative values indicate faster AV responses relative to the corresponding unimodal condition whereas positive values indicate a unimodal advantage. Asterisks mark significant within-participant differences between auditory and visual indices (paired t-tests, BH–FDR corrected, n = 42).

Relative RT measures showed that AV responses were substantially faster than auditory-only responses across conditions. For coherent stimuli, AV responses were 24–29% faster than auditory-only trials. Under non-coherent contexts, AV responses remained 7–15% faster than auditory-only performance depending on congruence. AV-congruent trials consistently yielded larger gains than AV-incongruent trials, and gains remained positive even in the most demanding blurred, non-coherent, incongruent condition (Fig. 5a).

### AV performance relative to the visual baseline

Relative to the visual baseline, the pattern depended more strongly on coherence and congruence (Supplementary Table 3). For coherent, clear AV-congruent stimuli, differences between AV and visual RTs were minimal. Visual blurring and non-coherent context increased AV gains relative to vision, yielding faster AV responses when the visual signal was degraded or contextually inconsistent. In contrast, AV-incongruent trials produced a clear AV cost relative to vision under coherent conditions. Visual blurring and non-coherent context reduced the magnitude of this incongruence-related cost but did not eliminate it. Notably, under non-coherent contexts, the advantage of AV congruency was reduced and expressed less consistently across conditions rather than uniformly abolished (Fig. 5b).

### Unimodal efficiency indices

Unimodal efficiency indices summarised AV performance relative to each unimodal stream (Fig. 5c). Indices relative to audition were consistently negative, indicating reliably faster AV than auditory-only responses. Indices relative to vision were generally near zero and became strongly positive mainly in coherent AV-incongruent trials, where AV responses were slower than visual-only responses. Paired comparisons showed that efficiency values relative to audition had larger magnitude than those relative to vision in nearly all conditions (all *q* ≤ 0.003), except clear non-coherent trials, where the two indices did not differ reliably (all *q* ≥ 0.17).

### Race Model Inequality: clarity, coherence, and congruence shape AV speed-up Race-model violations emerge as early deviations of AV RT distributions

To assess whether AV speed-ups exceeded what can be explained by independent processing of auditory and visual signals, we applied the race model inequality (RMI). Figure 6a illustrates the cumulative distribution functions (CDFs) of RTs for auditory-only, visual-only, and AV trials, plotted relative to each participant’s peak violation latency. The Miller bound defines the fastest responses expected under probability summation of independent auditory and visual channels; time points at which the AV CDF exceeds this bound indicate race-model violations, consistent with response-level coactivation.

**Figure 6.**
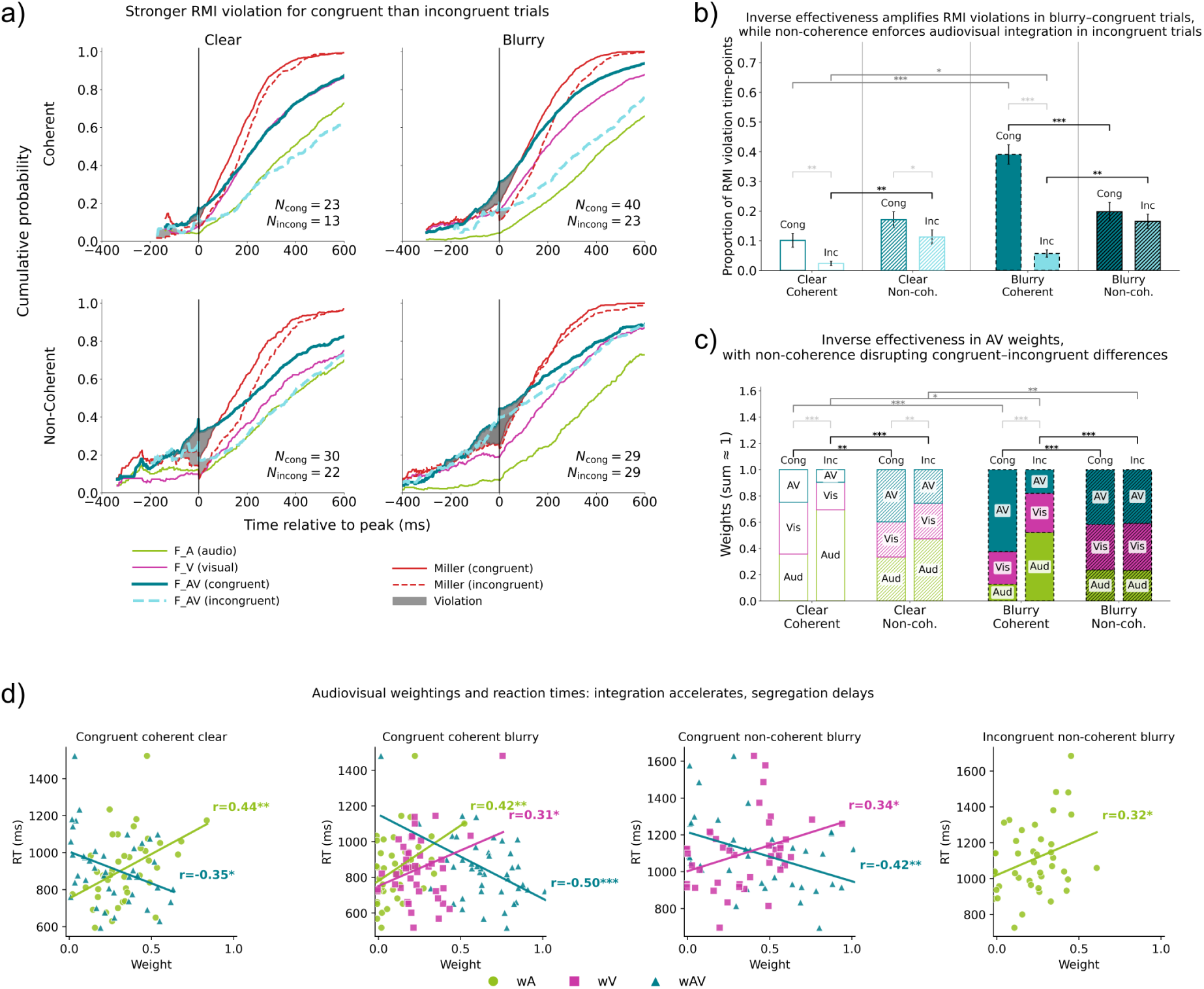
Multisensory response patterns vary with semantic coherence and cross-modal congruence. **(a)** Group-level cumulative RT distributions aligned to each participant’s peak Miller-violation latency. AV responses exceeded the Miller bound (shaded regions), with larger and earlier deviations observed for AV-congruent coherent conditions, particularly under blurring. Non-coherent conditions also exhibited elevated violation magnitudes across clarity levels. **(b)** Proportion of time points showing positive Miller violations (mean ± SEM). Blurring increased violation proportions for AV-congruent coherent conditions, whereas non-coherent conditions showed elevated values irrespective of congruency. Horizontal brackets denote significant within-subject contrasts surviving BH–FDR correction. **(c)** Descriptive sensory-weighting indices (wA, wV, wAV; constrained to sum to 1). Weight distributions differed across conditions, with more balanced contributions under clear coherent conditions and relatively higher auditory weights in coherent–incongruent conditions. Changes in visual clarity were associated with shifts in wAV across congruent and incongruent conditions. **(d)** Associations between response times and weighting indices. All uncorrected correlations are shown; only relations surviving BH–FDR correction are interpreted. Under blurry AV-congruent conditions, higher wAV values were associated with faster RTs, whereas under coherent conditions higher wA values corresponded to slower RTs. These relationships describe statistical correspondences between weighting indices and response latencies.

Across conditions, violations emerged as early leftward deviations of the AV CDF relative to the Miller bound, with their magnitude and temporal extent varying systematically with visual clarity, semantic coherence, and cross-modal congruence. Violations were most prominent for AV congruent trials, particularly when the visual signal was blurred, whereas incongruent AV trials showed smaller and more restricted deviations. Beyond group-level violation magnitudes, the prevalence of race-model violations across participants varied systematically across conditions (Fig. 6a). Under semantically coherent and AV-congruent conditions, the number of participants exhibiting violations increased markedly with visual degradation, rising from 23 of 42 participants in the clear condition to 40 of 42 participants in the blurry condition. This high prevalence indicates that when visual reliability was reduced but auditory and visual signals conveyed the same meaning, multisensory contributions to decision formation were engaged in the vast majority of participants. By contrast, although violations were also observed for AV-incongruent trials, their prevalence remained consistently lower than for congruent trials within each clarity level (clear: 13 vs. 23 participants; blurry: 23 vs. 40 participants), indicating that cross-modal conflict attenuated, but did not abolish, multisensory contributions. Under semantically non-coherent contexts, race-model violations were widespread across participants for both congruent and incongruent AV trials. Importantly, under blurry conditions, the number of participants exhibiting violations was identical for congruent and incongruent stimuli (29 of 42 participants in both cases; Fig. 6a). This convergence indicates that, when semantic information did not specify the upcoming concept, multisensory effects no longer depended on perceptual agreement between auditory and visual signals, but instead reflected convergence on the same behavioural response.

Importantly, violations of the race model inequality do not uniquely specify a perceptual integration mechanism. Such violations indicate that AV responses cannot be explained by independent parallel processing alone, but are compatible with multiple response-level architectures, including coactivation, correlated decision processes, or decisional pooling across modalities. Accordingly, RMI violations in the present study are interpreted as evidence for non-independent AV contributions to decision formation, rather than as evidence for perceptual fusion or a specific neural integration locus.

### Proportion of race-model violations

Quantifying these effects, the proportion of time points at which AV RT distributions exceeded the Miller bound varied reliably with visual clarity, semantic coherence, and AV congruence (Fig. 6b). Blurry targets produced a higher proportion of violation time points than clear targets (p < 0.001). Non-coherent primes yielded more violations than coherent primes (*p* = 0.028), and AV-incongruent trials produced fewer violations than AV-congruent trials (*p* = 0.014).

Interactions revealed that blurring selectively amplified violation proportions in coherent AV-congruent trials, with little or no increase in non-coherent or incongruent conditions (interaction ps < 0.005; Supplementary Table 4). A three-way interaction (*p* = 0.001) further indicated that, in non-coherent incongruent trials, blurring again increased violation proportions.

Variance partitioning (Supplementary Table 4) showed that visual clarity accounted for the largest share of unique variance in violation proportions (partial *R²* = 0.23), followed by congruence (0.16) and semantic coherence (0.10). Interactions together accounted for an additional partial *R²* of approximately 0.05–0.10. Overall, race-model violations were strongest when visual information was degraded but AV consistent, and were attenuated by cross-modal conflict.

### Peak violation latency reflects similar dependencies

Importantly, violations observed for AV incongruent targets occurred primarily under prime–target mismatch, where both modalities supported the same decision outcome despite encoding different concepts, indicating decision-level redundancy rather than perceptual congruence.

Peak RMI violation latency showed parallel dependencies on visual clarity, semantic coherence, and congruence (Fig. 6a). Each factor explained significant variance in peak latency (partial *R²* between 0.17 and 0.32), with additional contributions from their interactions (partial *R²* between 0.05 and 0.09). Non-coherent trials reached peak violation later than coherent trials (*p* = 0.006), whereas AV-incongruent trials reached peak violation earlier than AV-congruent trials (*p* = 0.020). Thus, both the magnitude and timing of AV speed-up depended jointly on sensory reliability and semantic context.

### Condition-specific weighting patterns

To characterize how auditory, visual, and integrative processes contributed to these RT distributions, we estimated descriptive weighting parameters for auditory (wₐ), visual (wᵥ), and combined (wₐᵥ) components (Fig. 6c).

In coherent, clear AV-congruent trials, weights were broadly balanced (wₐ = 0.358, wᵥ = 0.394, wₐᵥ = 0.248), with no significant pairwise differences after correction. In coherent, clear AV-incongruent trials, auditory weighting increased markedly (wₐ = 0.693), while the combined component decreased (wₐᵥ = 0.096), and wₐ exceeded both wᵥ and wₐᵥ (Supplementary Table 5).

In non-coherent AV-congruent trials, the combined component tended to dominate in both clear and blurry conditions, although these contrasts did not survive correction. Visual blurring had its strongest effect in coherent AV-congruent trials, where wₐᵥ increased to 0.624 and exceeded both wᵥ and wₐ (all corrected *p*s < 0.05). In coherent AV-incongruent blurry trials, auditory weighting remained dominant, whereas in non-coherent AV-incongruent blurry trials, visual and combined components exceeded auditory weighting, with only the wᵥ–wₐ contrast surviving correction (Supplementary Table 5).

### Associations between weighting parameters and RTs

Finally, correlation analyses between weighting parameters and RTs revealed condition-specific relationships (Fig. 6d). After correction, stronger auditory weighting was correlated with longer RTs in AV-congruent trials, both for coherent and blurry conditions. In contrast, higher combined weighting was correlated with shorter RTs in AV-congruent coherent blurry trials and AV-congruent non-coherent blurry trials. These correlations indicate that greater integrative weighting was selectively linked to faster responses under visually degraded conditions (Supplementary Table 6).

### Two RT profiles with comparable overall speed

An exploratory clustering of normalized, condition-wise RT profiles identified two participant subgroups (Group 1: *n* = 15; Group 2: *n* = 27; Fig. 7a). Clustering was performed on full RT profiles rather than on multisensory gain or cost indices, avoiding circularity with subsequent analyses. Overall mean RTs were comparable between groups (Group 1: 1067 ms; Group 2: 1089 ms), indicating that subgroup differences reflected condition-specific response patterns rather than general response speed. All subgroup analyses were exploratory and intended to characterize variability in multisensory behaviour rather than to define discrete participant types.

**Figure 7.**
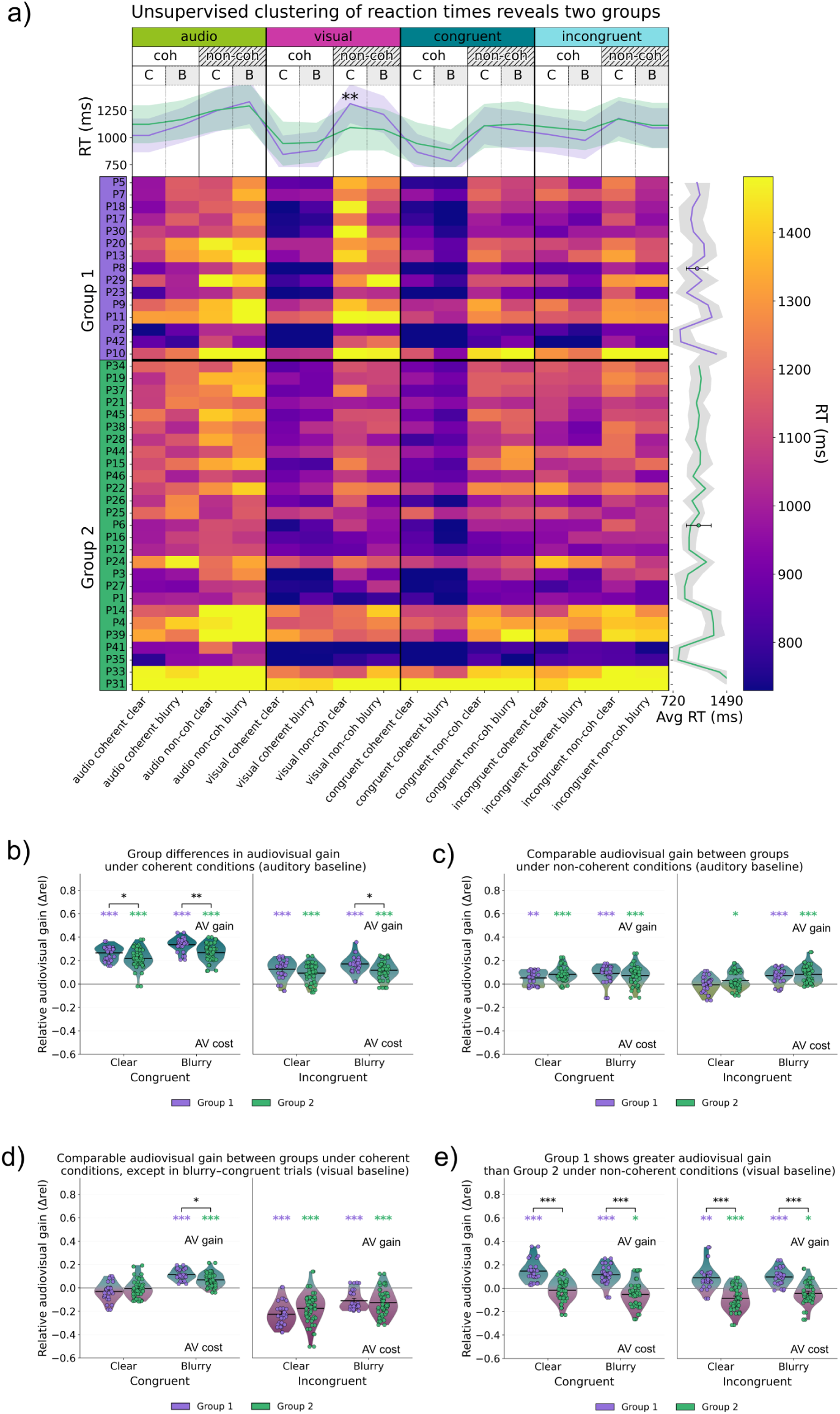
Exploratory RT clustering identifies two participant groups with distinct gain patterns. (a) Exploratory, data-driven clustering of condition-wise RT profiles. A participant-by-condition heatmap shows mean RTs (ms) across all modality × coherence × clarity combinations. Participants are ordered according to an unsupervised k-means solution (Group 1, purple; Group 2, green) computed on normalized RT profiles. Within each group, rows are sorted by Euclidean distance to the corresponding centroid in the original RT space. Columns correspond to unimodal auditory, unimodal visual, AV-congruent, and AV-incongruent conditions, further separated by semantic coherence (coherent vs. non-coherent) and visual clarity (clear vs. blurry). Smoothed group-average RT profiles shown above the heatmap summarize condition-wise differences across groups. (b) Relative AV gain computed using the auditory-only condition as baseline (Δ = [RT_audio − RT_AV] / RT_audio) for coherent trials. Under these conditions, Group 1 exhibited larger auditory-baseline AV gains than Group 2, particularly under blurry vision, as indicated by significant post-hoc contrasts. (c) Relative auditory-baseline AV gain for non-coherent trials. AV gains were reduced overall and did not differ reliably between groups across clarity levels. (d) Relative AV gain computed using the visual-only condition as baseline (Δ = [RT_visual − RT_AV] / RT_visual) for coherent trials. Visual-baseline gains were near zero for AV-congruent trials and strongly negative for AV-incongruent trials in both groups, with limited group differences. (e) Relative visual-baseline AV gain for non-coherent trials. Under these conditions, Group 1 showed consistently larger AV gains than Group 2 across congruency and clarity levels, most prominently for blurry incongruent trials, where AV responses were faster relative to the corresponding visual-only RTs.

### Condition-wise RT differences

Across conditions, the two groups exhibited partially overlapping but systematically different RT profiles (Fig. 7a). Under coherent contexts, Group 2 tended to respond more slowly than Group 1 across both clear and blurry conditions, particularly in AV trials. For example, in coherent clear AV-congruent trials, RTs averaged 761 ms in Group 1 and 847 ms in Group 2, with a similar separation observed for coherent AV-incongruent trials (887 ms vs. 973 ms). Differences of comparable magnitude were also present in coherent unimodal conditions.

In contrast, the largest between-group differences emerged under non-coherent visual conditions, where the pattern reversed. In the clear non-coherent visual-only condition, Group 1 responded substantially more slowly than Group 2 (1312 ms vs. 1092 ms, p < 0.01), with a similar difference observed under blurry non-coherent vision (1213 ms vs. 1074 ms). Under non-coherent contexts, Group 1 often produced AV RTs that were similar to or longer than visual-only RTs, particularly when the visual signal was clear, whereas Group 2 showed comparatively faster visual responses. These opposing tendencies under coherent versus non-coherent contexts motivated a closer examination of multisensory gains relative to unimodal baselines.

### Auditory-baseline multisensory gains by subgroup

When AV performance was evaluated relative to the auditory baseline (Fig. 7b,c), both groups showed clear multisensory gains under coherent contexts, indicating faster responses to AV than to auditory-only stimuli. However, the magnitude of this auditory-baseline facilitation differed by subgroup. In coherent conditions, Group 1 exhibited larger relative gains than Group 2, particularly under blurry vision, where between-group differences survived correction. Under non-coherent contexts, auditory-baseline gains were smaller overall and more variable, with no reliable group differences across clarity levels. Thus, subgroup differences in auditory-baseline facilitation were most evident when semantic expectations were aligned and auditory information benefitted from the presence of a visual signal.

### Visual-baseline multisensory gains by subgroup

A complementary pattern emerged when AV performance was evaluated relative to the visual baseline (Fig. 7d,e). Here, subgroup differences were selective to non-coherent contexts. Under semantic non-coherence, Group 1 showed positive visual-baseline gains, indicating faster AV than visual-only responses, whereas Group 2 showed negative gains, reflecting slower AV responses relative to vision. This divergence was observed for both congruent and incongruent trials and at both clarity levels, yielding reliable group differences after correction (all *q* ≤ 0.013). Averaged across non-coherent conditions, Group 1 exhibited modest but consistent gains (means approximately 0.08–0.16), whereas Group 2 showed corresponding costs (means approximately −0.12 to −0.02).

In contrast, under coherent contexts, visual-baseline gains were near zero in both groups for AV-congruent trials and strongly negative for AV-incongruent trials, with no reliable group differences. Thus, subgroup differences in visual-baseline multisensory effects emerged specifically when the prime–target relationship didn’t satisfy the semantic expectations.

Together, these results reveal complementary subgroup-dependent patterns across auditory and visual baselines. Group 1 expressed stronger auditory-baseline facilitation under coherent contexts and retained AV advantages relative to vision under non-coherent contexts. Group 2, by contrast, showed weaker auditory-baseline gains under coherence and a consistent visual advantage under non-coherent contexts, reflected in AV slowing relative to vision. These patterns indicate that individuals differed not in overall speed, but in how they leveraged AV information depending on semantic context and the unimodal reference frame.

### Exploratory subgroup differences in RMI, weighting, and subtitle use Group differences in violation proportions across clarity, coherence, and congruency

Both groups showed higher proportions of positive race-model inequality (RMI) violations under visual blurring and prime–target coherence, with the largest values observed for AV-congruent trials (Fig. 8a-c). Four conditions produced significant between-group differences after correction. In the clear non-coherent AV-congruent condition, Group 1 showed a higher violation proportion (0.171) than Group 2 (0.113; *p* < 0.004) (Fig. 8c). A group difference was also present in the clear non-coherent AV-incongruent condition (*p* = 0.038). Under blurry vision, violation proportions increased overall. In blurry coherent AV-congruent trials, Group 1 again showed higher values (0.391; *p* = 0.035). In blurry non-coherent trials, Group 1 showed higher violation proportions in both AV-congruent (0.198; *p* < 0.005) and AV-incongruent (0.165; *p* < 0.005) conditions. Together, these effects indicate that between-group differences in RMI magnitude were most consistently observed under non-coherent and visually degraded contexts. Inspection of individual-level CDFs further revealed that subgroup differences reflected not only variation in violation magnitude, but also differences in how consistently race-model violations were expressed across participants (Fig. 8a–b). Under non-coherent conditions, violations were observed in a larger proportion of participants in Group 1 than in Group 2 across both clarity levels. For example, in the clear non-coherent AV-congruent condition, violations were present in 13 of 15 participants in Group 1, compared with 17 of 27 participants in Group 2. Under blurry non-coherent conditions, violations were observed in most Group 1 participants (14–15 participants across congruency levels), whereas they were present in a smaller subset of Group 2 participants (15–25 participants depending on congruency).

**Figure 8.**
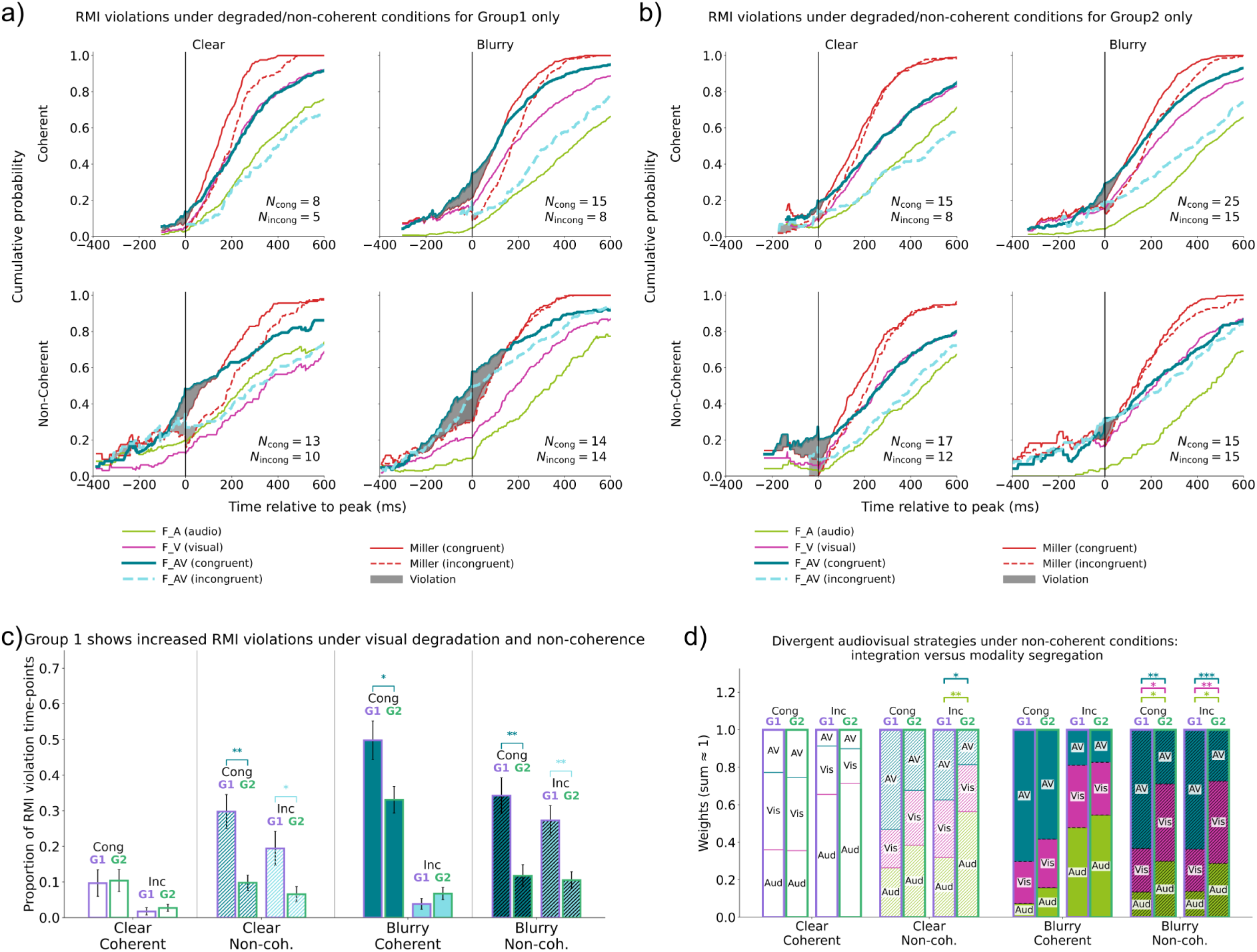
Group-specific patterns of race-model inequality violations and multisensory weighting. (a) (a) Cumulative distribution functions (CDFs) for Group 1, plotted by visual clarity (clear, blurry) and semantic context (coherent, non-coherent). Unimodal auditory, unimodal visual, and AV (congruent, incongruent) CDFs are shown together with the Miller bound (black line). Shaded regions indicate time points at which the AV CDF exceeds the bound, corresponding to RMI violations. (b) Corresponding CDFs for Group 2, displayed using an identical layout to facilitate comparison of violation timing and magnitude across groups. (c) Proportion of time points exhibiting positive RMI violations (mean ± SEM) for each clarity × coherence × congruence condition. Bars show group means with overlaid individual participant values; horizontal brackets denote conditions with significant between-group differences. (d) Descriptive AV weighting profiles for each group, decomposed into auditory (wA), visual (wV), and integrative (wAV) components. Stacked bars represent condition-wise estimated marginal means (constrained to sum to 1), with brackets indicating significant between-group differences across clarity and coherence contexts.

### Group differences in peak violation latency

Mean latencies of maximal RMI violation ranged from approximately 630 to 985 ms across conditions. Latencies were shorter in coherent trials and longer in non-coherent trials under blurry vision, but no between-group differences survived correction (all corrected *p*s ≥ 0.31 in clear conditions and ≥ 0.92 in blurry conditions). Overall, no reliable subgroup differences were observed in the timing of peak RMI violations.

For auditory weighting (wₐ), Group 2 showed higher values in several non-coherent conditions. In clear non-coherent AV-incongruent trials, wₐ was larger in Group 2 than in Group 1 (0.560 vs. 0.319; *p* = 0.00027), with additional differences in blurry non-coherent AV-congruent (*p* = 0.0246) and AV-incongruent (*p* = 0.0279) trials. For visual weighting (wᵥ), Group 2 showed higher values in blurry non-coherent conditions, with significant contrasts in AV-congruent (*p* = 0.0155) and AV-incongruent (*p* = 0.0058) trials. No significant between-group differences in wᵥ were observed in clear coherent or blurry coherent conditions.

### Group differences in weighting parameters

Race-model weighting parameters (wₐ, wᵥ, and wₐᵥ) showed systematic variation across groups and conditions (Fig. 8d). Subgroup differences were most pronounced for the combined component wₐᵥ in non-coherent contexts. In blurry non-coherent AV-congruent trials, wₐᵥ was higher in Group 1 (0.634) than in Group 2 (0.295; p = 1.89 × 10⁻⁷). A similar difference was observed in blurry non-coherent AV-incongruent trials (0.638 vs. 0.279; *p* = 5.85 × 10⁻⁸). Significant group differences in wₐᵥ were also present in clear non-coherent AV-congruent (p = 0.0019) and AV-incongruent (*p* = 0.0049) trials, whereas no differences emerged in coherent clear conditions.

### Exploratory links to self-reported viewing habits

An exploratory examination of the demographic questionnaire suggested a parallel between the two behavioural profiles and self-reported subtitle use. One question asked whether participants regularly use subtitles when watching movies. In Group 1, 11 participants responded “yes” and 2 responded “no” (84.6% “yes”). An exact binomial test confirmed that this proportion was significantly higher than a 50/50 distribution (*p* = 0.024), with a large associated effect size (Cohen’s *h* = 1.06). In Group 2, responses were evenly split (14 “yes,” 13 “no”). These exploratory findings indicate that the two RT-based subgroups also differed in their reported use of subtitles, suggesting a possible correspondence between viewing habits and the behavioural profiles observed in this task.

## Discussion

Multisensory facilitation is often described as a behavioural consequence of stimulus redundancy, modulated by sensory reliability and attenuated by cross-modal conflict (Ernst & Bülthoff, 2004; Stein & Stanford, 2007; Otto et al., 2013; Rohe & Noppeney, 2018). In everyday situations, however, people must decide what is happening based on sensory signals that vary in reliability, may conflict across modalities, and are interpreted relative to expectations about what is likely to occur. Despite extensive work on multisensory facilitation, it has remained unclear how sensory reliability, semantic expectations, and cross-modal correspondence jointly shape decisions about dynamic, naturalistic events, and why multisensory availability sometimes improves performance, sometimes has little effect, and sometimes incurs measurable costs. The present task required participants to evaluate explicit prime–target correspondence, thereby engaging semantic access, expectation matching, and decision-level evidence evaluation rather than low-level perceptual fusion. Importantly, these outcomes also vary substantially across individuals, raising the question of whether variability primarily reflects noise or a general integration ability, or instead reflects systematic strategy differences that emerge under specific decision constraints (Mahoney et al., 2011; Magnotti & Beauchamp, 2018). In the present task, the prime was designed to establish an expectation at the level of meaning rather than to preactivate specific sensory features. Prior work indicates that such primes activate conceptual representations in semantic memory, increasing the accessibility of task-relevant meanings and biasing interpretation and response selection (Bar, 2007; Doehrmann & Naumer, 2008; de Lange et al., 2018). This semantic framing provides a unifying account of the present results: it explains why multisensory effects generalized across alternative exemplars, remained robust under repetition, and supported response-level convergence even when auditory and visual signals were perceptually incongruent.

The present study shows that behavioural facilitation can increase both when expectations are met and when they are violated, but through distinct mechanisms revealed by race-model analyses. When semantic expectations were correct and auditory and visual signals were congruent, degrading the dominant sensory modality increased the contribution of the complementary modality, leading to faster responses and violations of the race-model inequality. In this context, reduced reliability promoted the joint use of congruent sensory information supported by an accurate prediction. Crucially, behavioural facilitation also emerged when predictions did not align with the incoming evidence. Here, race-model violations indicated joint multisensory contributions at the level of the response, even when auditory and visual signals were incongruent. Multisensory information was therefore recruited to resolve the mismatch, rather than being suppressed by audiovisual conflict. These results suggest an adaptive decision process that adjusts multisensory information use depending on predictive success or failure, consistent with broader accounts of perception as an interaction between incoming sensory evidence and prior expectations (Friston, 2005; Clark, 2013; de Lange et al., 2018). We showed that (1) when expectations guide perception effectively, multisensory use enhances efficiency under reduced sensory reliability, and (2) when expectations are not met, multisensory contributions support the resolution of unexpected events. Importantly, individuals differed in how they implemented this adaptive strategy, with some relying on joint multisensory use and others maintaining modality-specific processing under uncertainty.

### Semantic coherence determines when multisensory availability facilitates or impairs decisions

A central contribution of this work is demonstrating that multisensory availability alone is insufficient to predict behavioural facilitation. Instead, facilitation emerged selectively when semantic coherence was present between the prime and the target and when auditory and visual information jointly supported that judgement. In fact, at the group level, performance was highest when the prime and target were semantically coherent and audiovisual signals were congruent (Fig. 3). These conditions yielded the highest accuracy, the fastest responses, and the most compact RT distributions (Figs. 3, 4e), reflecting efficient decision formation when semantic context concorded with the expected response and both sensory streams provided converging evidence.

By contrast, when audiovisual signals were incongruent, multisensory input imposed a behavioural cost relative to visual-only performance (Figs. 4d, 5b). In these trials, the prime biased participants toward a specific judgement, one sensory modality supported that judgement, and the other opposed it, introducing direct competition during decision formation. This pattern aligns with prior demonstrations that semantic compatibility and cross-modal conflict jointly shape audiovisual behaviour (Doehrmann & Naumer, 2008; Chen & Spence, 2010; Gau & Noppeney, 2016).

### Sensory reliability modulates multisensory contributions at the level of response dynamics

Sensory reliability is a well-established factor influencing multisensory behaviour, often discussed in terms of inverse effectiveness, whereby multisensory benefits become more apparent when one sensory channel is degraded (Ernst & Bülthoff, 2004; Stein & Meredith, 1993; Holmes, 2007). In the present task, reducing visual clarity did not uniformly slow responses across conditions. Instead, its effect depended on both modality and semantic coherence, such that interactions between these factors explained RT patterns better than any main effect alone (Fig. 4d). Although multisensory behaviour is often discussed in terms of reliability-based weighting (Fetsch et al., 2011), the present results show that sensory reliability interacts with semantic expectations to shape how sensory information contributes to responses.

When visual information was clear, visual-only performance was often sufficient to resolve the semantic judgement, and AV responses closely matched visual RTs in semantically coherent and congruent contexts (Fig. 5b). Under these conditions, adding auditory information provided little behavioural advantage relative to vision. The relatively poor efficiency of audition in the present task likely reflects the nature of the decision rather than a general auditory disadvantage. Participants were required to map brief dynamic events onto abstract semantic categories and to evaluate prime–target correspondence, a process strongly supported by visual scene information. Because the prime itself was auditory, target sounds may have competed with the retained auditory representation of the prime, reducing the efficiency of audition when presented alone and rendering it the least effective unimodal pathway for this semantic judgement (Fig. 5a, c). However, when visual reliability was reduced, this balance changed. Importantly, inverse effectiveness effects must be interpreted relative to baseline variability and statistical structure (Holmes, 2009), a point addressed here by evaluating audiovisual performance relative to both auditory and visual reference frames. In semantically coherent and AV-congruent trials, degraded visual input increased the behavioural relevance of auditory information, such that combined AV evidence could surpass visual-only performance (Figs. 4d–e, 5b). Thus, reduced sensory reliability did not enhance multisensory contributions indiscriminately. Rather, it increased the likelihood that both auditory and visual information influenced the response, specifically in contexts where they conveyed consistent information about whether the target matched the prime.

This interaction was most clearly expressed at the level of RT distributions. Under blurred, semantically coherent, and AV-congruent conditions, RT distributions showed a selective speed-up in their fastest quantiles (Fig. 4e), indicating earlier response commitment rather than a uniform shift across the distribution. These distributional effects provide information beyond mean RTs by revealing when multisensory contributions influence decision dynamics rather than later processing stages.

Beyond their temporal profile, it is also important to determine whether these multisensory effects were stimulus-specific or generalized across different instances of the same semantic category. Each concept was therefore presented twice using distinct visual realizations (e.g., different videos depicting the same event category), preserving conceptual identity while varying low-level visual form and contextual details. The persistence of the main effects across first and second presentations indicates that the patterns of facilitation, cost, and race-model violation were not driven by idiosyncratic stimulus features or one-off associations, but reflected stable mappings between semantic expectations, sensory information, and decision processes. Although repetition produced modest overall speed-ups, consistent with limited practice effects reported in short experimental paradigms (Duff et al., 2012), these gains did not qualitatively alter the structure of multisensory interactions. Instead, multisensory benefits and costs remained robust across repeated encounters with the same concept, suggesting that the effects observed here reflect stable decision-level dynamics rather than transient learning or training effects tied to specific trials or exemplars.

### Semantic non-coherence reveals heterogeneous strategies for using multisensory information

When the prime did not match the target, semantic information was present but did not provide a useful prediction about the upcoming event. In these non-coherent trials, performance costs were observed across conditions, but multisensory behaviour became markedly more variable across participants (Fig. 5). Crucially, semantic non-coherence did not abolish multisensory effects. Instead, it reduced the functional distinction between audiovisual congruent and incongruent events. In these contexts, neither modality aligned with the prime-based expectation, such that congruent audiovisual information no longer provided a clear advantage, and incongruent information did not impose an additional cost. As a result, congruent and incongruent conditions became more similar in their behavioural impact, relative to semantically coherent contexts (Fig. 5b). This pattern explains why, under non-coherence, audiovisual congruency did not strongly differentiate performance despite clear effects under coherent conditions. Rather than reflecting a uniform reduction in multisensory processing, semantic non-coherence reveals heterogeneous decision strategies, with individuals differing in the extent to which they rely on combined versus modality-specific information under uncertainty.

### Race-model evidence reveals decision-level convergence beyond probability summation

Race-model analyses further clarify the nature of multisensory contributions observed in this task. Traditionally, race models have been applied to redundant target paradigms to distinguish probability summation from joint contributions of multiple sensory signals (Miller, 1982; Otto & Mamassian, 2012; Innes & Otto, 2019). Under the race model, auditory and visual signals are processed independently and in parallel, with the faster channel triggering the response. This framework defines an upper limit on facilitation expected from probability summation alone, known as the Miller bound.

In the present study, race-model inequality violations were not a general feature of audiovisual stimulation. Instead, violation proportions increased selectively when reduced visual reliability coincided with semantic coherence and audiovisual congruence (Fig. 6b), with early violation latencies. These effects indicate departures from independent processing specifically in contexts where combining auditory and visual information improved behavioural efficiency. Converging descriptive evidence came from weighting parameters, which showed increased combined audiovisual contributions under the same conditions (Fig. 6c).

Importantly, the present findings extend the use of race-model logic beyond classic redundant target designs. Here, race-model evidence tracked convergence at the level of the decision outcome required by the task, rather than perceptual congruence between auditory and visual signals. In semantically non-coherent contexts, race-model violations were observed even for audiovisual-incongruent events (Fig. 6b). In these trials, auditory and visual signals differed in content but both supported the same decision, namely that the target did not match the prime. This demonstrates that race-model signatures can arise when multiple sensory signals jointly support a decision outcome, even in the absence of perceptual agreement between modalities, in line with prior demonstrations that multisensory speed-ups can reflect non-independent decision processes rather than perceptual fusion (Miller, 1982; Otto & Mamassian, 2012; Innes & Otto, 2019; Chua et al., 2022). Together, these results indicate that race-model violations can arise through at least two functionally distinct routes: (1) increased multisensory contributions when visual information is degraded and when AV signals jointly support the same interpretation, and (2) response-level convergence when auditory and visual information both favor the same decision outcome despite conveying different content. Importantly, these effects do not imply a single underlying computational architecture.

Correlations between reaction times and weighting parameters provided additional support for this interpretation. Across several conditions, higher combined audiovisual weighting was associated with faster responses, whereas higher visual and auditory weighting was associated with slower responses (Fig. 6d). Importantly, these relationships were observed in both semantically coherent and non-coherent contexts, and were strongest under conditions of increased uncertainty, including visual degradation and semantic non-coherence. Although these associations are descriptive rather than mechanistic, their convergence with distribution-level speed-ups and race-model violations supports the interpretation that contexts in which combined sensory information contributes more strongly are also those in which behavioural efficiency improves.

### Individual differences reflect strategy differences when semantic information does not allow reliable anticipation of the target

Exploratory clustering revealed two participant subgroups with comparable overall response speed but systematically different multisensory decision profiles (Fig. 7a). Importantly, all trials contributing to these analyses were correct responses; group differences therefore do not reflect failures in task performance, but differences in how sensory information was used to reach a decision. Differences between groups were minimal when semantic coherence between the prime and the target provided a reliable prediction about the upcoming event (Figs. 7b–e). By contrast, when the prime did not match the target, decisions could not be prepared in advance on the basis of semantic correspondence, and clear differences emerged in how participants used the available auditory and visual information to reach a response.

One subgroup showed larger audiovisual advantages relative to vision, stronger race-model inequality violations, and higher combined audiovisual weighting in semantically non-coherent contexts (Figs. 7–8). This pattern suggests a strategy in which participants seek cross-modal confirmation when semantic predictions do not indicate which sensory stream should be prioritised. By contrast, the other subgroup showed smaller audiovisual advantages and more unimodal weighting, indicating stronger independent reliance on each modality in the same context. These differences do not reflect reduced perceptual efficiency, but distinct adaptations to decisional uncertainty revealed specifically when semantic information did not allow reliable anticipation of the upcoming concept. This is consistent with prior evidence that individual variability in multisensory behaviour reflects stable information-processing strategies rather than a unitary integration ability (Mahoney et al., 2011; Magnotti & Beauchamp, 2018; Otto et al., 2013).

This interpretation is further supported by the observation that the subgroup relying more strongly on combined information showed selective slowing in unimodal visual trials under semantic non-coherence (Fig. 8), particularly when visual input was clear. In these cases, visual information alone was sufficient to reach the correct decision, but the absence of a reliable semantic prediction appeared to delay commitment in individuals who typically rely on multisensory confirmation. When auditory information was unavailable, this strategy could not be implemented, resulting in slower responses despite intact visual evidence.

The exploratory association between behavioural profiles and self-reported subtitle use provides a tentative link between laboratory measures of multisensory decision strategies and everyday audiovisual experience. Regular subtitle use may be associated with a greater tendency to rely on concurrent visual and linguistic cues to stabilise meaning, particularly when auditory information is uncertain or insufficient on its own. Although preliminary, the subtitle-use pattern is in line with the broader view that experience can calibrate how observers weight concurrent cues. Together, these findings indicate that individual variability in multisensory behaviour reflects systematic differences in decision strategy that become evident when semantic correspondence does not allow advance anticipation of the target, rather than differences in perceptual ability or response accuracy.

### Implications, broader relevance, and limitations

Taken together, the present findings identify two distinct routes by which multisensory information contributes to behavioural responses, demonstrating that facilitation depends on how meaning and prior expectations guide the use of sensory information, rather than arising as a fixed consequence of stimulus redundancy. This framework explains why redundant-signal speed-ups emerged in some contexts but not others, and why evidence for non-independence was selectively expressed when reduced sensory reliability and semantic alignment encouraged joint contributions from auditory and visual information. Crucially, the present results also indicate that multisensory contributions can arise through a different route when semantic expectations are not met. In these contexts, cross-modal incongruency did not map directly onto reduced multisensory effects, as convergence could still emerge when auditory and visual signals supported the same response, even in the absence of perceptual agreement between modalities. Importantly, these two modes of multisensory contribution differed not only in their contextual triggers, but also in how broadly they were expressed across participants. When visual reliability was reduced under semantically coherent and AV-congruent conditions, race-model violations were observed in the vast majority of participants, consistent with a reliability-driven shift toward joint multisensory contributions during response generation. By contrast, under semantic mismatch, violations were widespread but less uniformly expressed across individuals, consistent with a more flexible, strategy-dependent mode in which perceptually distinct signals converged on the same response. Notably, the identical prevalence of violations for congruent and incongruent audiovisual events under semantic non-coherence provides strong evidence that multisensory effects in this context reflect response-level convergence rather than shared perceptual content. When both auditory and visual signals supported the same judgement, perceptual correspondence no longer differentiated behaviour. Together, these prevalence patterns indicate that the two routes identified here reflect distinct processes of multisensory response use, rather than graded variations of a single integrative mechanism.

Beyond multisensory response times, the present findings have broader implications for understanding how perception, attention, and judgment interact in everyday cognition (Bar, 2007; Talsma et al., 2010; Choi et al., 2018; de Lange et al., 2018). The strong dependence of multisensory effects on semantic expectations suggests that perceptual decisions are shaped by access to conceptual memory, consistent with evidence that semantic knowledge constrains interpretation early in decision formation (Bar, 2007; Ellis et al., 2015). Moreover, the selective engagement of multisensory information when it supports a decision goal aligns with views of attention as a mechanism for prioritizing task-relevant evidence under uncertainty rather than uniformly enhancing sensory signals (Talsma et al., 2010; Choi et al., 2018).

From a comprehensive decision-making perspective, the present results help explain why expectations can both facilitate perception and introduce systematic biases in real-world judgments. When multiple information sources converge on the same categorical response, decisions are accelerated; when expectations bias interpretation toward a specific outcome, conflicting evidence can instead slow or distort responses. Such dynamics are directly relevant to situations such as eyewitness judgments, rapid event interpretation, and reasoning under uncertainty, where expectations and sensory evidence interact to shape confidence and accuracy rather than perception alone. At a broader level, these findings align with evidence that multisensory interactions unfold across multiple temporal and decisional scales, with later stages reflecting context- and task-dependent convergence rather than early sensory fusion (Senkowski et al., 2011; Senkowski & Engel, 2024).

Several considerations delimit the scope of these conclusions. First, the task probed multisensory behaviour in an explicit semantic correspondence judgement and therefore primarily reflects decision-level contributions rather than early perceptual fusion. This framing is not a limitation per se, but an important boundary on interpretation. Second, although the stimuli were dynamic and naturalistic, the semantic space was necessarily constrained for experimental control, which may limit generalisation to more open-ended or graded semantic environments. Third, modality asymmetries and multisensory gains should be interpreted relative to task demands and decision requirements, rather than as modality-invariant properties of audiovisual processing. Fourth, subgroup analyses were exploratory and require replication across tasks and validation using independent behavioural or neural measures. Finally, the sample consisted of young, French-speaking adults from a relatively homogeneous cultural background, and semantic associations and multisensory strategies may vary across populations.

Future work extending this paradigm to broader semantic spaces that vary across languages and cultures (Buchanan et al., 2025), and to cross-task validation, will be essential for clarifying how multisensory decision strategies adapt to sensory reliability, semantic expectations, and individual experience. In parallel, combining this paradigm with neural measures will be critical for testing whether the two behavioural routes identified here are supported by distinct neural dynamics or by flexible recruitment of shared decision-related circuits.

## Methods

### Participants

Forty-two adults (18–40 years; *M* = 24.3, *SD* = 4.8) participated in the study. Participants were recruited through online platforms and reported normal or corrected-to-normal vision and normal hearing. Individuals with a history of neurological disorders or unmanaged psychiatric conditions were excluded. All participants provided informed consent in accordance with institutional ethical guidelines and received monetary compensation for approximately 45 minutes of participation.

Participants completed a brief demographic questionnaire including age and gender, as well as exploratory questions concerning everyday multisensory habits (e.g., subtitle use and perceived sensory reliance). Questionnaire data were not used to define experimental conditions and were analysed for exploratory purposes only.

### Stimuli

The experiment followed a semantic correspondence paradigm in which each trial began with an auditory prime naming a semantic concept, followed by a unimodal or AV target stimulus, and a binary coherence judgment (Fig. 1a).

The stimulus set comprised 117 naturalistic AV events drawn from five semantic categories: humans, animals, objects, transport, and environments (Fig. 1b). Category membership was balanced across the experiment and treated as a controlled design dimension rather than an analytical factor.

Each AV stimulus consisted of a short video clip paired with a temporally aligned sound. AV pairings were either congruent, with auditory and visual components originating from the same semantic event, or incongruent, with auditory and visual components drawn from different semantic categories (Fig. 1a).

A repetition manipulation was included such that semantic concepts were encountered twice using different visual exemplars. Repetition preserved conceptual identity while varying low-level visual input, allowing semantic reactivation to be assessed independently of visual familiarity (Fig. 1a).

Visual reliability was manipulated by generating blurred versions of the videos using Microsoft Clipchamp. The blur operation reduced high spatial-frequency information while preserving global scene structure (Fig. 1c). Only the visual component was altered; auditory recordings were identical across clarity conditions.

### Experimental design

Stimuli were presented across four blocks that varied in visual clarity and exemplar repetition. Participants encountered both clear and blurry stimuli as well as first and second exemplars of the same semantic concepts. Block order was partially counterbalanced across participants to minimise confounding effects of time, fatigue, or practice. Trial order was randomised independently within each block.

Seven stimulus conditions were tested, comprising unimodal auditory trials, unimodal visual trials (clear and blurry), and AV trials that crossed visual clarity with cross-modal congruency (congruent, incongruent). This design established unimodal baselines and enabled multisensory benefits and costs to be quantified relative to both auditory and visual reference conditions under varying levels of visual reliability.

### Task and procedure

Participants were seated in an anechoic chamber approximately 60 cm (about 24 inches) from the computer monitor, such that their eyes were aligned with the center of the screen. Auditory stimuli were delivered directly through the computer’s speakers throughout the task. Each trial consisted of a spoken semantic prime (2.5 s), followed immediately by a target stimulus presented for 2 s in one of the experimental conditions.

Participants judged whether the target matched the preceding prime using a two-alternative forced-choice response (“match” vs “mismatch”). Response mapping was held constant across participants. Auditory stimuli were normalised in root-mean-square amplitude. The procedure was identical across blocks.

### Behavioural measures

RTs were measured from target onset to keypress. Trials were excluded if RTs were shorter than 150 ms, longer than 3000 ms, or exceeded 3.5 median absolute deviations within each participant and condition. Accuracy was recorded on each trial.

Multisensory facilitation or interference was quantified by comparing AV RTs with unimodal baselines using:

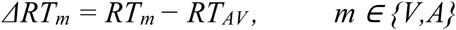

A positive value indicated that AV responses were faster than the corresponding unimodal baseline and therefore reflected a multisensory gain. A negative value indicated slower AV responses and therefore reflected a multisensory cost. These values were computed for every participant within each combination of clarity and congruency. They were used directly in the mixed-effects analyses, the race model evaluation, and the clustering procedure.

Rather than relying on composite indices such as inverse efficiency scores (Bruyer & Brysbaert, 2011), we analyzed accuracy and reaction times separately to avoid interpretational ambiguities when speed–accuracy trade-offs vary across conditions.

### Statistical analyses

An overview of the behavioural analyses is provided in Fig. 2. Analyses included accuracy and RTs, full RT distributions, race-model inequality metrics, descriptive sensory weighting indices, and exploratory clustering to characterize individual differences.

### Mixed-effects models

Accuracy was analysed using generalized linear mixed-effects models with a binomial logit link. Fixed effects included visual clarity, AV congruency, prime–target semantic coherence, and repetition. Random intercepts were specified for participants, with random slopes for clarity and congruency included when supported by model convergence. Statistical significance of fixed effects was assessed using likelihood ratio tests, and estimates are reported as odds ratios with corresponding confidence intervals.

RTs were analysed using linear mixed-effects models applied to log-transformed values. The fixed-effect structure matched that of the accuracy models. Models were fitted using restricted maximum likelihood. Effect sizes were summarised using semi-partial *R²* values. Planned contrasts tested the effects of AV incongruency, multisensory gains relative to unimodal baselines, and the interaction between visual degradation and congruency.

### Race Model Inequality (RMI)

We used a RMI framework to assess whether RTs in the AV condition could be explained by independent probability summation of the auditory and visual channels, or whether they instead reflected multisensory coactivation. The RMI provides an upper bound on the cumulative distribution function (CDF) of response probabilities to AV stimuli at a given latency, and violations of this bound indicate responses that are faster than expected under a simple race process, consistent with integrative processing (Mahoney & Verghese, 2019). To capture the structure of our task, the RMI was applied separately across conditions defined by AV congruency and visual clarity.

Formally, the race model predicts:

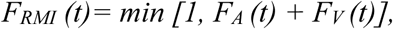

where *FA (t)* and *FV (t)* are the auditory and visual CDFs. We compared the observed AV CDF *FAV (t)* with this bound to determine whether multisensory responses exceeded what would be expected from independent processing of the two modalities.

For each subject and condition, CDFs were estimated for auditory, visual, and AV trials. Two standard RMI-derived metrics were computed: (i) the proportion of time points at which the AV CDF exceeded the Miller bound (violation proportion) and (ii) the latency of the peak violation (time of maximal AV–bound difference). Statistical inference was performed using within-subject bootstrap resampling (10,000 iterations) to generate null distributions for each metric under the assumption of no true coactivation. Group-level significance was assessed by comparing observed values with these nulls, and multiple comparisons across clarity conditions were controlled using a false discovery rate of q = 0.05.

Although race-model analyses are most commonly applied to redundant multisensory signals, their logic applies more generally to any condition in which multiple information sources can independently support the same behavioural decision. In the present task, the decision concerned semantic correspondence between the prime and the target. Under prime–target mismatch, both auditory and visual components of the target, even when conceptually incongruent with each other, could provide convergent evidence for a ‘mismatch’ response. Accordingly, race-model analyses were applied to both congruent and incongruent AV targets, with violations interpreted as evidence for non-independent contributions to decision formation rather than perceptual integration.

### Weight estimation based on RMI

To characterize the relative contributions of auditory, visual, and combined processes to RT distributions, a softmax mixture model was applied. For each participant and condition, three non-negative weights summing to one were estimated, corresponding to auditory-driven, visual-driven, and integrative response components.

Because the weights form compositional data, isometric log-ratio transformations were applied prior to statistical analysis. Linear mixed-effects models were fitted with clarity, congruency, and repetition as fixed effects and participant as a random effect. Planned contrasts examined changes in relative weighting across conditions.

These weights provide descriptive summaries of how response distributions reflect unisensory and combined influences and are not interpreted as estimates of perceptual or neural integration mechanisms. Weight patterns were consistent across bootstrap resamples and showed convergent but non-redundant relationships with race-model metrics, indicating that the weighting parameters capture complementary aspects of response-time structure rather than recapitulating RMI measures.

Correlations between weighting parameters, RTs, and RMI metrics were computed within conditions for descriptive purposes only. As all measures derive from the same RT distributions, these analyses are exploratory and not treated as independent tests.

### Unsupervised clustering on relative ΔRT

To characterize individual differences in multisensory behaviour, unsupervised clustering was performed on mean RTs across experimental conditions. Clustering was conducted on RTs rather than on multisensory gain or cost indices to avoid circularity with subsequent analyses.

RT values were standardised across participants prior to clustering. The two-cluster solution was supported by both elbow and silhouette criteria and remained stable across bootstrap resampling, supporting its use as a descriptive summary of individual differences rather than a definitive categorical classification. Clusters were used to describe variability in how participants responded to changes in visual reliability, semantic context, and AV congruency and were treated as exploratory descriptors rather than confirmatory groupings.

### Reporting and transparency

All statistical tests were two-sided. Multiple comparisons were controlled using false discovery rate correction where applicable. Data exclusion criteria were defined a priori and applied uniformly across conditions.

## Acknowledgment

This project was funded by the Canadian Institutes of Health Research (#166197, F.L., V.H., E.S., L.S-G.).

## Author contributions

E.S. contributed to the design of the experimental paradigm, led stimulus selection, participant recruitment, and data collection, performed initial analyses, and co-wrote the manuscript.

L.S.-G. contributed to stimulus selection and data collection.

K.J., S.B., and F.L. provided critical feedback on analyses, figures, and interpretation.

V.H. conceived the study, designed and programmed the experimental paradigm, performed the analyses and result visualizations, and co-wrote the manuscript.

## Corresponding author

Correspondence to Vanessa Hadid

## Annexe

**Table S1.** Mixed-effects model results and variance decomposition for task accuracy.

| Main effects |  |  |  |  |
| --- | --- | --- | --- | --- |
| Effect | Contrast | Accuracy (Mean ± SEM) |  | p-value |
| Modality | Congruent vs Incongruent vs Visual vs Audio | 0.97 vs 0.85 vs 0.96 vs 0.93 |  | < **0.001** |
| Coherence | Coherent vs Non-coherent | 0.95 vs 0.91 |  | **1.4 × 10<sup>-8</sup>** |
| Repetition | 2nd vs 1st | 0.95 vs 0.92 |  | **0.0016** |
| Clarity | Clear vs Blurry | 0.96 vs 0.94 |  | 0.78 |
| Interactions |  |  |  |  |
| Modality × Coherence × Clarity |  |  |  |  |
| Modality × Coherence | Clear (Mean ± SEM) | Blurry (Mean ± SEM) | Δ (Blurry – Clear) | q-value (BH–FDR) (Clear vs Blurry) |
| Audio · Coherent | 0.9211 ± 0.0062 | 0.9062 ± 0.0068 | −0.0149 | — |
| Audio · Non-coherent | 0.9467 ± 0.0127 | 0.9599 ± 0.0085 | +0.0132 | — |
| Visual · Coherent | 0.9514 ± 0.0064 | 0.9421 ± 0.0110 | −0.0094 | 0.405 |
| Visual · Non-coherent | 0.9393 ± 0.0134 | 0.9802 ± 0.0071 | +0.0409 | **0.00052** |
| AV-congruent · Coherent | 0.9824 ± 0.0033 | 0.9876 ± 0.0035 | +0.0052 | 0.378 |
| AV-congruent · Non-coherent | 0.9277 ± 0.0137 | 0.9802 ± 0.0082 | +0.0524 | **0.00073** |
| AV-incongruent · Coherent | 0.8023 ± 0.0131 | 0.8063 ± 0.0155 | +0.0039 | 0.793 |
| **Variance decomposition** |  |  |  |  |
| **Predictor** | **Unique variance explained (partial R<sup>2</sup>)** |  |  |  |
| Modality | 0.039 |  |  |  |
| Coherence | 0.018 |  |  |  |
| Repetition | 0.010 |  |  |  |
| Clarity | 0.005 |  |  |  |
| Modality × Coherence | 0.008 |  |  |  |

**Table S2.** Fixed-effect estimates from mixed-effects modeling and variance decomposition for RTs (log-RT).

| **Effect / Contrast** | **β ± SE (log-RT)** | **p-value** |
| --- | --- | --- |
| Modality (Audio vs Visual) | 0.173 ± 0.015 | **2.1 × 10<sup>-32</sup>** |
| Modality (Congruent vs Visual) | -0.003 ± 0.015 | 0.82 |
| Modality (Incongruent vs Visual) | 0.149 ± 0.018 | **3.5 × 10<sup>-17</sup>** |
| Semantic coherence (Non-coherent vs Coherent) | 0.270 ± 0.024 | **2.7 × 10<sup>-30</sup>** |
| Clarity (Blurry vs Clear) | 0.010 ± 0.016 | 0.58 |
| Repetition (2nd vs 1st) | -0.002 ± 0.019 | 0.95 |
| Audio × Blurry | 0.119 ± 0.021 | **1.9 × 10<sup>-8</sup>** |
| Congruent × Blurry | -0.094 ± 0.021 | **6.1 × 10<sup>-6</sup>** |
| Audio × Non-coherent | -0.144 ± 0.031 | **4.9 × 10<sup>-6</sup>** |
| Incongruent × Non-coherent | -0.149 ± 0.031 | **1.2 × 10<sup>-6</sup>** |
| Clarity × Coherence | -0.067 ± 0.029 | **0.022** |
| Congruent × Clarity × Coherence | 0.140 ± 0.040 | **5.4 × 10<sup>-4</sup>** |
| **Modality × Coherence × Clarity** |  |  |

| **Modality × Coherence** | **Clear (Mean ± SEM, ms)** | **Blurry (Mean ± SEM, ms)** | **Δ (Blurry – Clear, ms)** | **q-value (BH–FDR)** |
| --- | --- | --- | --- | --- |
| Audio · Coherent | 1085.5 ± 26.7 | 1154.4 ± 25.0 | +69.0 | **4.38 × 10<sup>-8</sup>** |
| Audio · Non-coherent | 1254.3 ± 29.5 | 1302.2 ± 31.2 | +47.9 | **5.30 × 10<sup>-4</sup>** |
| Visual · Coherent | 916.9 ± 31.2 | 925.1 ± 27.8 | +8.3 | 0.532 |
| Visual · Non-coherent | 1178.7 ± 35.2 | 1119.5 ± 31.0 | -59.2 | **0.0235** |
| Congruent · Coherent | 924.8 ± 30.0 | 845.6 ± 27.8 | -79.2 | **1.78 × 10<sup>-8</sup>** |
| Congruent · Non-coherent | 1115.8 ± 30.1 | 1098.0 ± 31.0 | -17.8 | 0.333 |
| Incongruent · Coherent | 1078.9 ± 28.8 | 1027.5 ± 27.5 | -51.4 | **2.84 × 10<sup>-3</sup>** |
| Incongruent · Non-coherent | 1182.5 ± 29.6 | 1099.3 ± 30.8 | -83.1 | **3.30 × 10<sup>-7</sup>** |
| **Variance decomposition (partial R<sup>2</sup>)** |  |  |  |  |
| **Predictor** | **Unique variance explained (partial R<sup>2</sup>)** |  |  |  |
| Modality | 0.1017 |  |  |  |
| Coherence | 0.0833 |  |  |  |
| Modality × Coherence | 0.0087 |  |  |  |
| Modality × Clarity | 0.0075 |  |  |  |
| Clarity | 0.0062 |  |  |  |
| Repetition | ~0 |  |  |  |

**Table S3.** Relative AV gain and cost with respect to unimodal baselines.

| **Baseline** | **Condition** | **t** | **p-value (uncorrected)** | **q-value (BH–FDR)** |
| --- | --- | --- | --- | --- |
| **Auditory** | Coherent · Clear · Congruent | -16.05 | **2.8 × 10<sup>-19</sup>** | **< 10<sup>-18</sup>** |
|  | Coherent · Blurry · Congruent | -18.57 | **1.4 × 10<sup>-21</sup>** | **< 10<sup>-20</sup>** |
|  | Coherent · Clear · Incongruent | -15.31 | **1.5 × 10<sup>-18</sup>** | **< 10<sup>-18</sup>** |
| | Coherent · Blurry · Incongruent | −17.14 | $2.6 \times 10^{-20}$ | $< 10^{-19}$ |
|  | Non-coherent · Clear · Congruent | −1.39 | 0.171 | n.s. |
|  | Non-coherent · Clear · Incongruent | −1.78 | 0.083 | n.s. |
| | Non-coherent · Blurry · Congruent | −4.04 | $2.26 \times 10^{-4}$ | $3.1 \times 10^{-4}$ |
| | Non-coherent · Blurry · Incongruent | −4.18 | $1.51 \times 10^{-4}$ | $2.4 \times 10^{-4}$ |
| **Visual** | Coherent · Clear · Congruent | −0.07 | 0.94 | n.s. |
| | Coherent · Clear · Incongruent | +10.44 | $4.1 \times 10^{-13}$ | $< 10^{-12}$ |
| | Coherent · Blurry · Congruent | −6.96 | $1.9 \times 10^{-8}$ | $7.6 \times 10^{-8}$ |
| | Coherent · Blurry · Incongruent | +13.93 | $3.9 \times 10^{-17}$ | $< 10^{-16}$ |
| | Non-coherent · Clear · Congruent | +5.36 | $3.5 \times 10^{-6}$ | $4.6 \times 10^{-6}$ |
| | Non-coherent · Clear · Incongruent | −5.36 | $3.5 \times 10^{-6}$ | $5.6 \times 10^{-6}$ |
|  | Non-coherent · Blurry · Congruent | +1.68 | 0.100 | n.s. |
|  | Non-coherent · Blurry · Incongruent | +0.08 | 0.937 | n.s. |

**Table S4.**
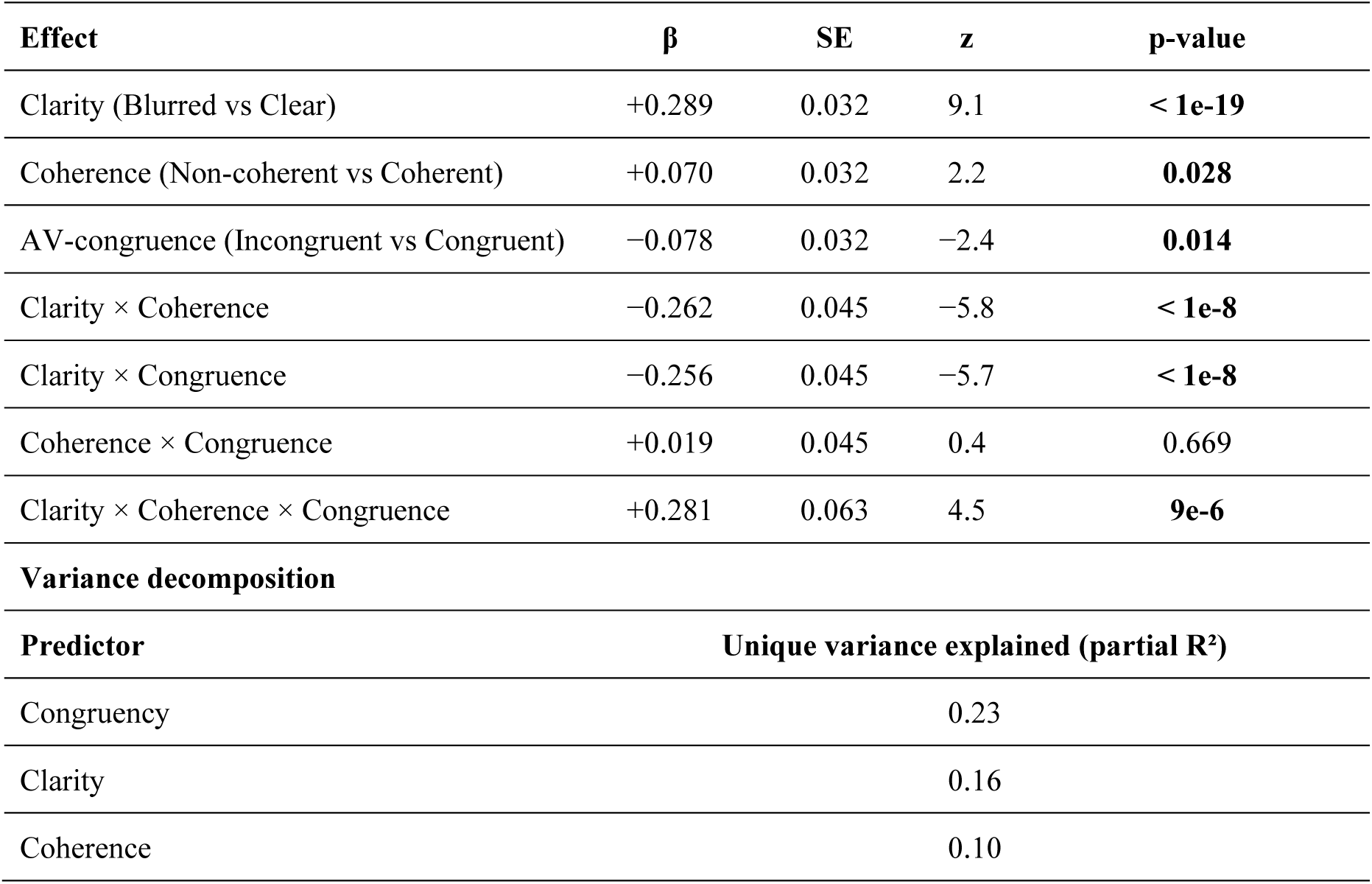
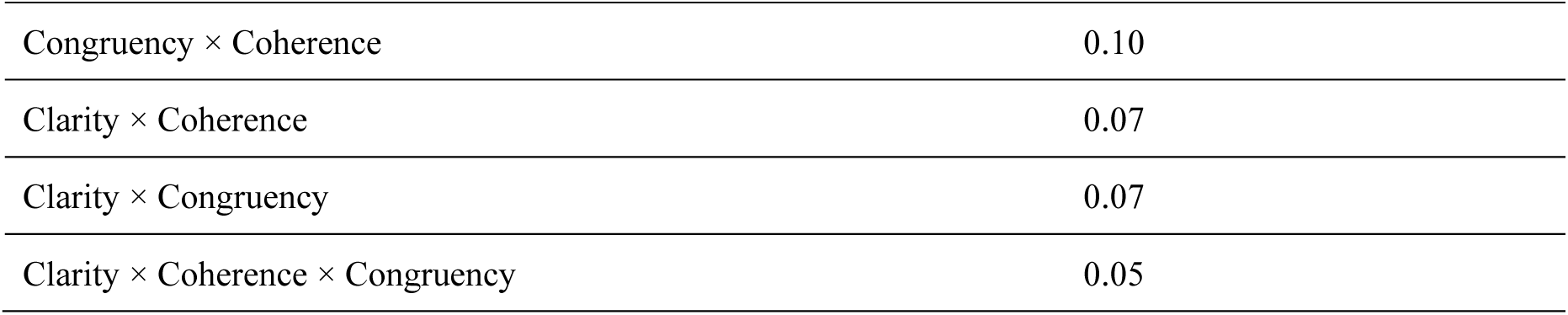
Effects of clarity, semantic coherence, and AV congruence on race-model violations.

| Effect | $\beta$ | SE | z | p-value |
| --- | --- | --- | --- | --- |
| Clarity (Blurred vs Clear) | +0.289 | 0.032 | 9.1 | $< 1e-19$ |
| Coherence (Non-coherent vs Coherent) | +0.070 | 0.032 | 2.2 | **0.028** |
| AV-congruence (Incongruent vs Congruent) | −0.078 | 0.032 | −2.4 | **0.014** |
| Clarity × Coherence | −0.262 | 0.045 | −5.8 | $< 1e-8$ |
| Clarity × Congruence | −0.256 | 0.045 | −5.7 | $< 1e-8$ |
| Coherence × Congruence | +0.019 | 0.045 | 0.4 | 0.669 |
| Clarity × Coherence × Congruence | +0.281 | 0.063 | 4.5 | **9e-6** |
| **Variance decomposition** |  |  |  |  |
| Predictor | Unique variance explained (partial R<sup>2</sup>) |  |  |  |
| Congruency | 0.23 |  |  |  |
| Clarity | 0.16 |  |  |  |
| Coherence | 0.10 |  |  |  |

|  |  |
| --- | --- |
| Congruency × Coherence | 0.10 |
| Clarity × Coherence | 0.07 |
| Clarity × Congruency | 0.07 |
| Clarity × Coherence × Congruency | 0.05 |

**Table S5.**
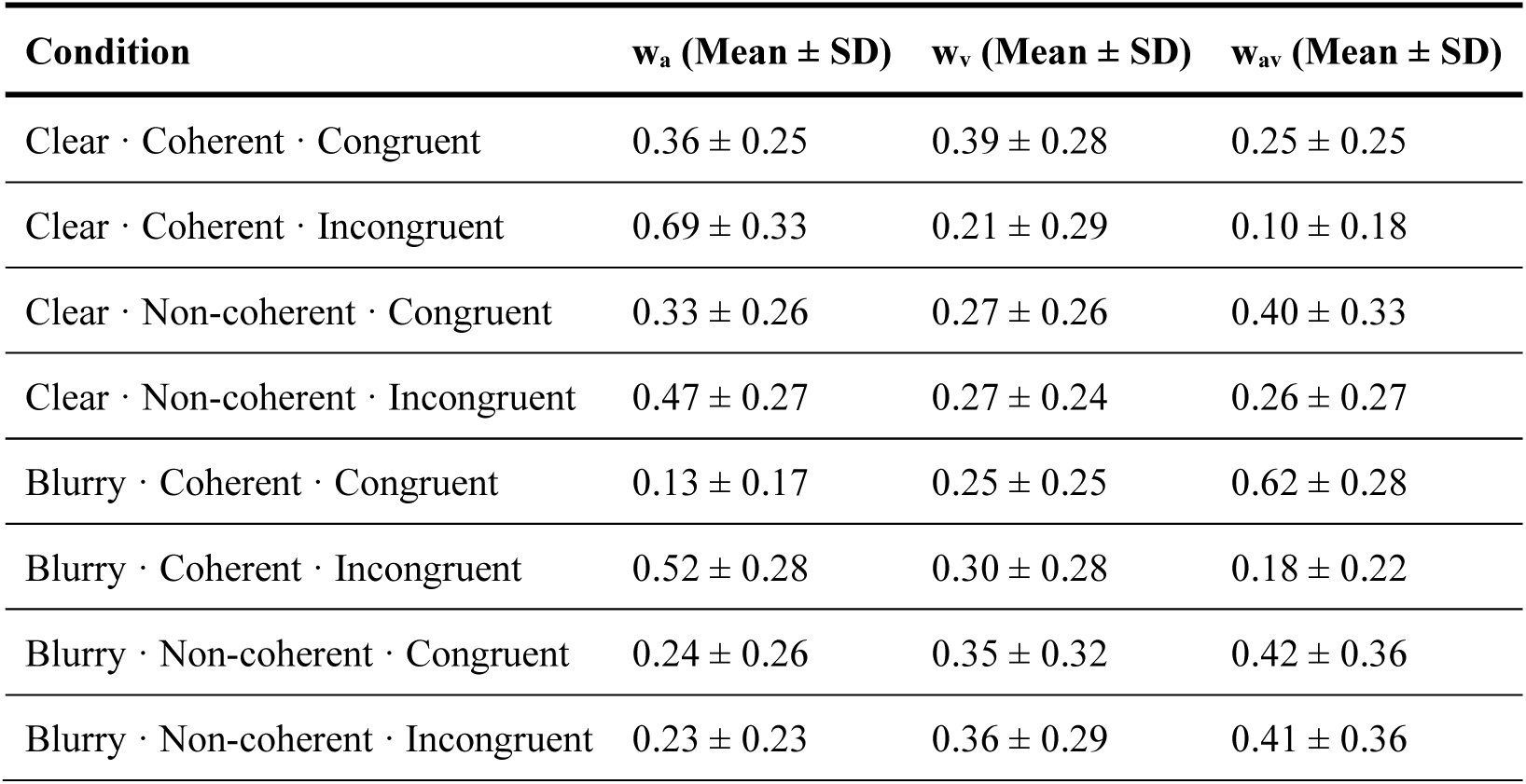
Condition-specific weighting of auditory, visual, and integrative components.

| Condition | $w_a$ (Mean ± SD) | $w_v$ (Mean ± SD) | $w_{av}$ (Mean ± SD) |
| --- | --- | --- | --- |
| Clear · Coherent · Congruent | 0.36 ± 0.25 | 0.39 ± 0.28 | 0.25 ± 0.25 |
| Clear · Coherent · Incongruent | 0.69 ± 0.33 | 0.21 ± 0.29 | 0.10 ± 0.18 |
| Clear · Non-coherent · Congruent | 0.33 ± 0.26 | 0.27 ± 0.26 | 0.40 ± 0.33 |
| Clear · Non-coherent · Incongruent | 0.47 ± 0.27 | 0.27 ± 0.24 | 0.26 ± 0.27 |
| Blurry · Coherent · Congruent | 0.13 ± 0.17 | 0.25 ± 0.25 | 0.62 ± 0.28 |
| Blurry · Coherent · Incongruent | 0.52 ± 0.28 | 0.30 ± 0.28 | 0.18 ± 0.22 |
| Blurry · Non-coherent · Congruent | 0.24 ± 0.26 | 0.35 ± 0.32 | 0.42 ± 0.36 |
| Blurry · Non-coherent · Incongruent | 0.23 ± 0.23 | 0.36 ± 0.29 | 0.41 ± 0.36 |

**Table S6.** Condition-specific correlations between model-derived weighting parameters and RTs.

| Weight | Condition | $r$ | p-value (uncorrected) | q-value (BH-FDR) |
| --- | --- | --- | --- | --- |
| $w_A$ | Congruent · Coherent · Clear | 0.438 | **0.003** | **0.035** |
| $w_A$ | Incongruent · Coherent · Clear | −0.020 | 0.899 | 0.899 |
| $w_A$ | Congruent · Non-coherent · Clear | 0.267 | 0.088 | 0.234 |
| $w_A$ | Incongruent · Non-coherent · Clear | −0.039 | 0.807 | 0.877 |
| $w_A$ | Congruent · Coherent · Blurry | 0.422 | **0.005** | **0.035** |
| $w_A$ | Incongruent · Coherent · Blurry | 0.238 | 0.129 | 0.287 |
| $w_A$ | Congruent · Non-coherent · blurry | 0.236 | 0.132 | 0.287 |
| $w_A$ | Incongruent · Non-coherent · Blurry | 0.324 | **0.036** | 0.124 |
| $w_V$ | Congruent · Coherent · Clear | −0.034 | 0.832 | 0.877 |
| $w_V$ | Incongruent · Coherent · Clear | −0.095 | 0.549 | 0.693 |
| wV | Congruent · Non-coherent · Clear | −0.032 | 0.841 | 0.877 |
| wV | Incongruent · Non-coherent · Clear | 0.177 | 0.263 | 0.485 |
| wV | Congruent · Coherent · Blurry | 0.314 | **0.042** | 0.127 |
| wV | Incongruent · Coherent · Blurry | −0.117 | 0.461 | 0.650 |
| wV | Congruent · Non-coherent · Blurry | 0.341 | **0.027** | 0.108 |
| wV | Incongruent · Non-coherent · Blurry | −0.057 | 0.720 | 0.864 |
| wAV | Congruent · Coherent · Clear | −0.352 | **0.022** | 0.105 |
| wAV | Incongruent · Coherent · Clear | 0.152 | 0.338 | 0.506 |
| wAV | Congruent · Non-coherent · Clear | −0.215 | 0.171 | 0.342 |
| wAV | Incongruent · Non-coherent · Clear | −0.109 | 0.493 | 0.657 |
| wAV | Congruent · Coherent · Blurry | −0.502 | **0.001** | **0.017** |
| wAV | Incongruent · Coherent · Blurry | −0.158 | 0.316 | 0.505 |
| wAV | Congruent · Non-coherent · Blurry | −0.418 | **0.006** | **0.035** |
| wAV | Incongruent · Non-coherent · Blurry | −0.160 | 0.313 | 0.505 |

